# Single-cell RNA sequencing reveals distinct tumor microenvironmental patterns in lung adenocarcinoma

**DOI:** 10.1101/2020.12.11.419606

**Authors:** Philip Bischoff, Alexandra Trinks, Benedikt Obermayer, Jan Patrick Pett, Annika Lehmann, Philipp Jurmeister, Aron Elsner, Tomasz Dziodzio, Jens-Carsten Rückert, Jens Neudecker, Christine Falk, Dieter Beule, Christine Sers, Markus Morkel, David Horst, Nils Blüthgen, Frederick Klauschen

## Abstract

Recent developments in immuno-oncology demonstrate that not only cancer cells, but also features of the tumor microenvironment guide precision medicine. Still, the relationship between tumor and microenvironment remains poorly understood. To overcome this limitation and identify clinically relevant microenvironmental and cancer features, we applied single-cell RNA sequencing to lung adenocarcinomas. While the highly heterogeneous carcinoma cell transcriptomes reflected histological grade and activity of relevant oncogenic pathways, our analysis revealed two distinct microenvironmental patterns. We identified a prognostically unfavorable group of tumors with a microenvironment composed of cancer-associated myofibroblasts, exhausted CD8+ T cells, proinflammatory monocyte-derived macrophages and plasmacytoid dendritic cells (CEP^2^ pattern) and a prognostically favorable group characterized by myeloid dendritic cells, anti-inflammatory monocyte-derived macrophages, normal-like myofibroblasts, NK cells and conventional T cells (MAN^2^C pattern). Our results show that single-cell gene expression profiling allows to identify patient subgroups based on the tumor microenvironment beyond cancer cell-centric profiling.

## Introduction

Lung cancer accounts for the majority of new cases and deaths of cancer worldwide [1]. The most common subtype of lung cancer is lung adenocarcinoma, which is at the same time the most prevalent subtype in non-smokers [2]. Although some patients can be cured by surgery complemented with radiotherapy or chemotherapy, the overall prognosis of lung cancer is still poor. Molecular profiling identified a variety of driver mutations and guides treatment with novel targeted therapies, such as inhibitors of epidermal growth factor receptor (EGFR) or anaplastic lymphoma kinase (ALK) [3, 4]. Moreover, expression of PD-L1 in tumor cells predicts response to the recently introduced class of immune checkpoint inhibitors [5]. However, clinical data shows that the predictive accuracy of PD-L1 expression alone is limited pointing to more complex mechanisms in the relationship between tumor cells and their microenvironment. These findings suggest that the classical focus on the tumor cells is not capturing the full picture. A more comprehensive characterization including the tumor microenvironment is likely required for a better understanding of tumor biology and more precise patient stratifications.

Cancers are multicellular communities comprising malignant epithelial cells and different types of non-malignant immune and stromal cells which exhibit dynamic and reciprocal interactions. Modulation of immune responses, remodeling of the extracellular matrix and neoangionesis essentially determine the aggressiveness of cancer [6]. In vitro and xenograft studies outline relevant roles for different cell types of the tumor microenvironment. Current bulk omics analyses do not allow high-resolution in situ characterization of tumors uncovering the cellular diversity of the tumor microenvironment. However, comprehensive profiling of tumor microenvironmental cell types and their functional traits in patient tissue is essential to estimate the clinical relevance of results from preclinical models.

In this study, we analyzed tumor epithelial cells and associated non-malignant cells of the tumor microenvironment of lung adenocarcinomas. We utilized tissue samples from routine lung surgery to generate single-cell libraries using a commercial droplet-based approach and investigated gene expression profiles on the single-cell level. This way, we demonstrate high interpatient heterogeneity in gene expression of tumor epithelial cells while histological subtypes still share common transcriptomic traits. Within the tumor microenvironment, we identified various immune and stromal cell subtypes, of which many were depleted or enriched in tumor tissue. Extending previous single-cell studies [7–10], we found that the heterogeneous cellular composition of the tumor microenvironment across patients follows specific patterns that were associated with the differentiation grade of carcinoma cells. We evaluated these signatures in an independent retrospective cohort characterized by bulk gene expression profiling and found that these patterns might correlate with patient prognosis. We conclude that a comprehensive profiling of lung adenocarcinomas considering the cellular diversity of tumor-associated non-malignant cells may help reveal novel tumor subtypes based on carcinoma cells but also microenvironmental features, and provide clinically relevant information guiding novel therapies targeting components of the tumor microenvironment.

## Results

### Single-cell RNA sequencing uncovers the cellular diversity of lung adenocarcinomas

To study the cellular composition of lung adenocarcinomas with a particular focus on the microenvironment on the single-cell level, 10 normal lung and 10 lung adenocarcinoma fresh tissue samples were collected during routine lung surgery. Evaluation of consecutive H&E stained tissue sections ensured proper sampling of tumor tissue and showed that tumor morphology ranged from well differentiated lepidic to poorly differentiated sarcomatoid growth patterns (Supp. Fig. 1). Tissue samples were enzymatically dissociated to a single-cell suspension to produce single-cell transcriptome libraries using a commercial droplet-based system (Fig. 1A). Unsorted single-cell RNA sequencing yielded in 133,736 single-cell transcriptomes of which 114,489 high-quality transcriptomes remained after quality control and filtering (for quality control parameters see Supp. Fig. 2).

**Figure 1:**
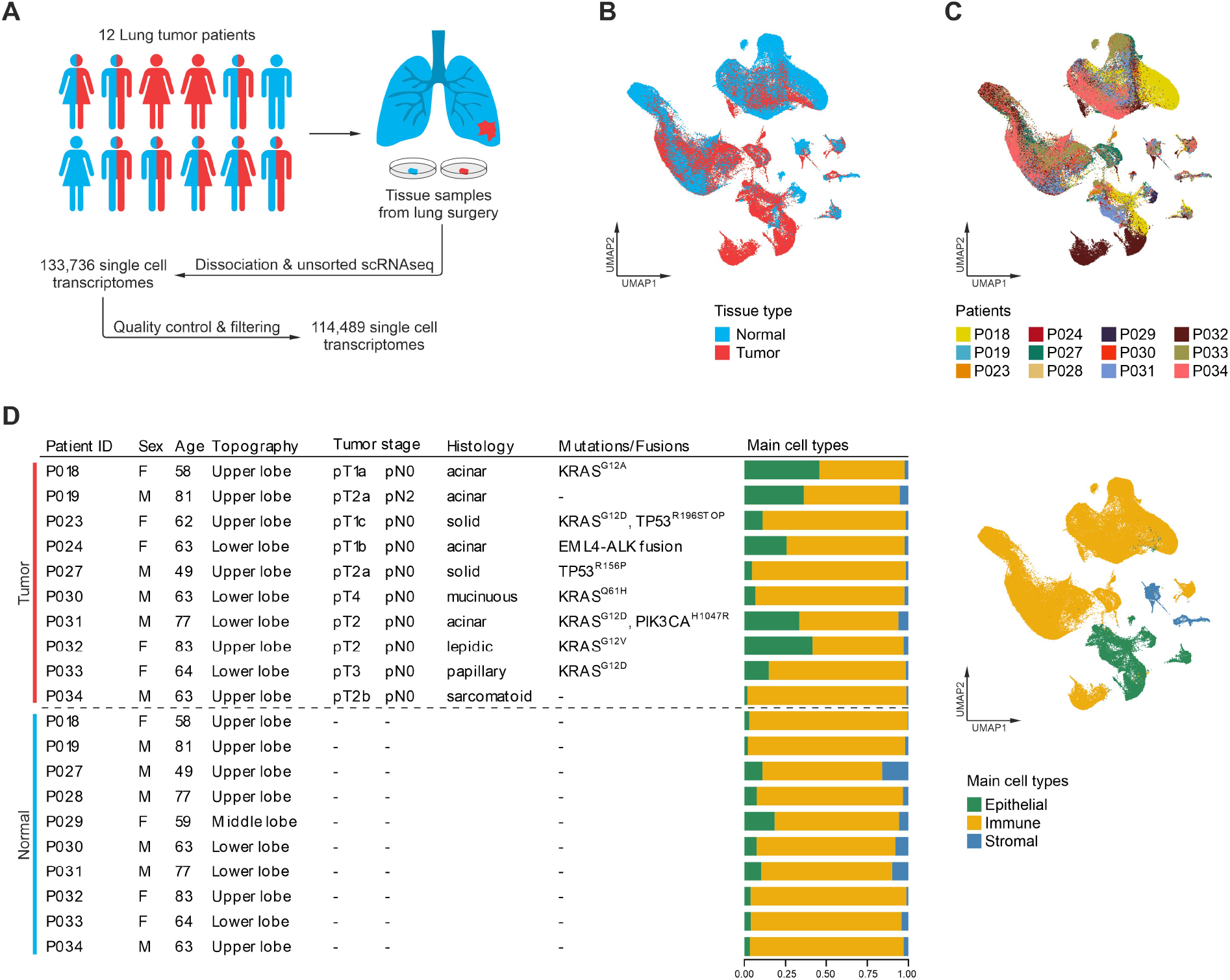
Single-cell RNA sequencing of lung adenocarcinomas. (A) Schematic representation of the workflow, 10 normal (blue) and 10 tumor (red) tissue samples were obtained from 12 patients. (B+C) UMAPs based on the top 15 principal components of all single-cell transcriptomes after filtering, color-coded by (B) tissue type, or (C) patient. (D) Overview of clinical features, clinically relevant oncogenic mutations and gene fusions; quantification of main cell types per patient and UMAP of all single-cell transcriptomes color-coded by main cell type.

Analysis and visualization by Uniform Manifold Approximation and Projection (UMAP) [11] showed that single-cell transcriptomes of different tissue types or patients both intermingled in many clusters, excluding general batch effects, and partly formed tumor- or patient-specific clusters, indicating underlying biological differences (Fig. 1B-C). To uncover which cellular compartments account most for interpatient variability, we analyzed single-cell transcriptomes for the expression of epithelial, immune and stromal marker genes. In total, 20,450 epithelial, 89,766 immune, and 4,273 stromal single-cell transcriptomes were covered, suggesting an overrepresentation of immune cell transcriptomes as observed in other studies [9, 12]. The yield of epithelial transcriptomes varied depending on histological subtypes, as we observed less than 10% in solid/sarcomatoid, but up to >40% in lepidic/acinar carcinomas (Fig. 1D). Epithelial cells showed the highest degrees of interpatient heterogeneity (Fig. 1C-D).

### Intertumoral heterogeneity of tumor epithelial cells reflects differentiation gradients

To further dissect interpatient variability within the epithelial cell compartment, epithelial single-cell transcriptomes were subset and reclustered (Supp. Fig. 3A). Epithelial clusters overrepresented in normal or tumor tissue samples were defined as normal or tumor cell clusters, respectively (Supp. Fig. 3B). Within the normal cell clusters, we found alveolar type 1 and 2, club, ciliated, and even a small cluster of neuroendocrine cells (Fig. 2A), which were characterized by expression of typical individual marker genes (Fig. 2B) and gene signatures (Supp. Fig. 4A-B) [13, 14]. The club cell cluster also expressed basal cell marker genes such as *NGFR* and *KRT5* indicating an admixture of small amounts of basal cells in this cluster (Fig. 2B). Few single-cell transcriptomes of normal epithelial cell types were also obtained from tumor tissue samples (Fig. 2A), probably due to normal tissue adjacent to or entrapped within the tumor. Beyond that, tumor cell clusters distinctly segregated from the normal cell clusters (Fig. 2A). In contrast to normal cell clusters, the tumor cell clusters were mainly patient-specific, indicating high intertumoral heterogeneity of tumor epithelial cells (Fig. 2A). This was underlined by a variety of genes differentially expressed across tumors such as *EGFR, TFF3, CDKN2A,* and *SFTPA2* (Fig. 2C, black arrowheads), correlating with protein expression as shown by immune staining (Fig. 2D). In order to analyze functional intertumoral heterogeneity, we inferred the activity of potentially oncogenic signaling pathways from single-cell gene expression profiles [15, 16]. We found highly varying pathway activity scores for EGFR, TGFβ, JAK/STAT, Hypoxia, and PI3K signaling in tumor epithelial cells of different patients (Fig. 2E), and these patterns were largely unrelated to the mitotic activity of tumor epithelial cells (Supp. Fig. 3D). The activity of p53 signaling was significantly reduced in tumors harboring *TP53* mutations, demonstrating the functional effects of genomic alterations. In contrast, pathway activity scores for EGFR and MAPK signaling were not significantly higher in *KRAS*-mutated compared to *KRAS*-wildtype tumors, suggesting that other mechanisms can activate MAPK in cancer (Supp. Fig. 3C). Taken together, tumor epithelial cells exhibited high intertumoral heterogeneity on the levels of gene expression, protein expression and signaling pathway activity.

**Figure 2:**
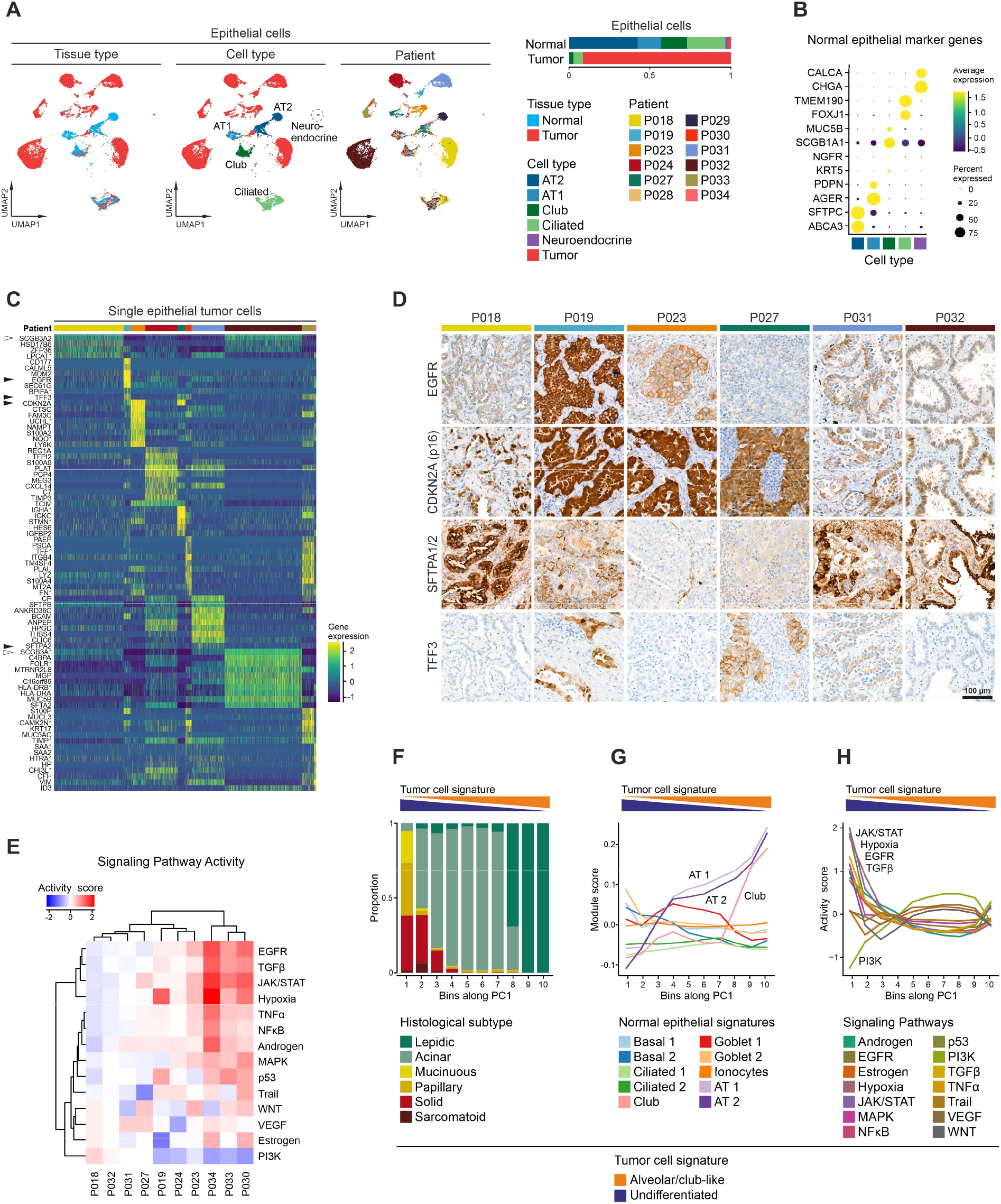
Intertumoral heterogeneity of tumor epithelial cells in lung adenocarcinomas. (A) UMAPs based on the top 20 principal components of all epithelial single-cell transcriptomes color-coded by tissue type, cell type and patient, and quantification of epithelial cell types per tissue type, AT1 = alveolar type 1 cells, AT2 = alveolar type 2 cells. (B) Average gene expression of selected marker genes for normal epithelial cell types. (C) Differentially expressed genes in tumor epithelial cells grouped by patients, maximum top 10 genes showed per patient, for patient color code see (A). (D) Immunohistochemical staining of proteins encoded by selected differentially expressed genes indicated by black arrowheads in (C). (E) Mean pathway activity scores of tumor epithelial cells grouped by patient. (F) Distribution of histological subtypes, (G) mean module scores of normal epithelial cell type gene signatures, and (H) mean pathway activity scores of tumor epithelial cells sorted along principal component 1 (PC1). (F+G+H) Principal component analysis based on gene expression of all tumor epithelial single-cell transcriptomes; schematic depiction of tumor cell signature module scores along PC1.

Despite obvious intertumoral heterogeneity of tumor epithelial cells, we also noted shared features of tumor transcriptomes across patients. Some differentially expressed genes were strongly expressed in more than one tumor cases such as *SCGB3A1* in patients P018, P031 and P032, and *SCGB3A2* in P018 and P032 (Fig. 2C, white arrowheads). Additionally, tumors shared signaling pathway activity patterns. Pathway activity scores for EGFR, TGFβ and JAK/STAT signaling were high in tumors of patients P030, P033 and P034, while the highest PI3K signaling activity was found in P018 and P032 (Fig. 2E). In order to emphasize similarities between tumors from different patients, we performed UMAP embedding based on fewer principal components (4, 6, 8 instead of 20). In low-dimensional UMAPs, tumor cells clustered less by patient, but rather by histological subtypes (Supp. Fig. 3E). Here, the first principal component (PC1) displayed a gradient from poorly and moderately (sarcomatoid, solid, papillary, mucinuous) to moderately (acinar) to well differentiated tumors (lepidic) (Fig. 2F, Supp. Fig. 3E+G). Interestingly, *SCGB3A1* and *SCGB3A2*, two genes that were previously associated with lung development [17], positively correlated with this dimension (Supp. Fig. 3F, arrowheads). Moreover, gene signature scores of normal lung cell types [13] along PC1 showed a strong positive correlation with gene expression profiles of alveolar type 1 and 2 as well as club cells (Fig. 2G). Together, this indicates that PC1 reflects the degree of differentiation of tumor epithelial cells. Hence, the top 30 genes positively and negatively correlated with PC1 were defined as an “alveolar/club-like” and “undifferentiated” tumor cell signature, respectively (Fig. 2F-H, Supp. Fig. 3I).

To further characterize functional features of “alveolar/club-like” and “undifferentiated” tumor epithelial cells, we analyzed mitotic and pathway activity along PC1. While tumor cells with different degrees of differentiation exhibited no clear differences in mitotic activity (Supp. Fig. 3H), we found high pathway activity scores for JAK/STAT, Hypoxia, EGFR and TGFβ signaling in “undifferentiated”, and high scores for PI3K signaling in “alveolar/club cell-like” tumor epithelial cells, respectively (Fig. 2H). We conclude that tumor epithelial cells of different lung adenocarcinoma patients exhibit transcriptional patterns along a spectrum ranging from undifferentiated to differentiated alveolar/club cell-like phenotypes that correlate with distinct oncogenic pathway activity.

### Two subtypes of myofibroblasts constitute the tumor stromal microenvironment

Neovascularization and remodeling of the extracellular matrix impacts on tumor growth and metastasis. To analyze the cellular composition of the stromal microenvironment, we separately analyzed stromal cells from both normal and tumor tissue samples. We identified different clusters of endothelial and lymphatic endothelial cells, different clusters of fibroblasts, myofibroblasts and smooth muscle cells, as well as mesothelial cells (Fig. 3A, for normal tissue per patient see Supp. Fig. 5A) which were characterized by expression of typical marker genes (Fig. 3B) and gene signatures (Supp. Fig. 4A-B) [13, 14]. Among the endothelial cells clusters, tumor tissues were dominated by clusters 2 and 4 (Fig. 3A). These clusters showed high expression of genes involved in angiogenesis such as *VWA1* and *HSPG2*, as well as high expression of *INSR*, encoding a marker protein and possible therapeutic target in tumor-associated endothelial cells, consistent with a previous report [9] (Supp. Fig. 5B, arrowheads). While the proportion of smooth muscle and mesothelial cell clusters did not differ significantly between normal and tumor tissues, we detected a shift from fibroblast to myofibroblast cell clusters in tumor tissues (Fig. 3A). Here, myofibroblast clusters were characterized by expression of both fibroblastic marker genes, such as *PDGFRA* and *LUM*, and smooth muscle marker genes, such as *MYLK* and *ACTA2* (Fig. 3B). Hence, besides tumor-associated endothelial cells, myofibroblasts characterized the stromal microenvironment of lung adenocarcinomas.

**Figure 3:**
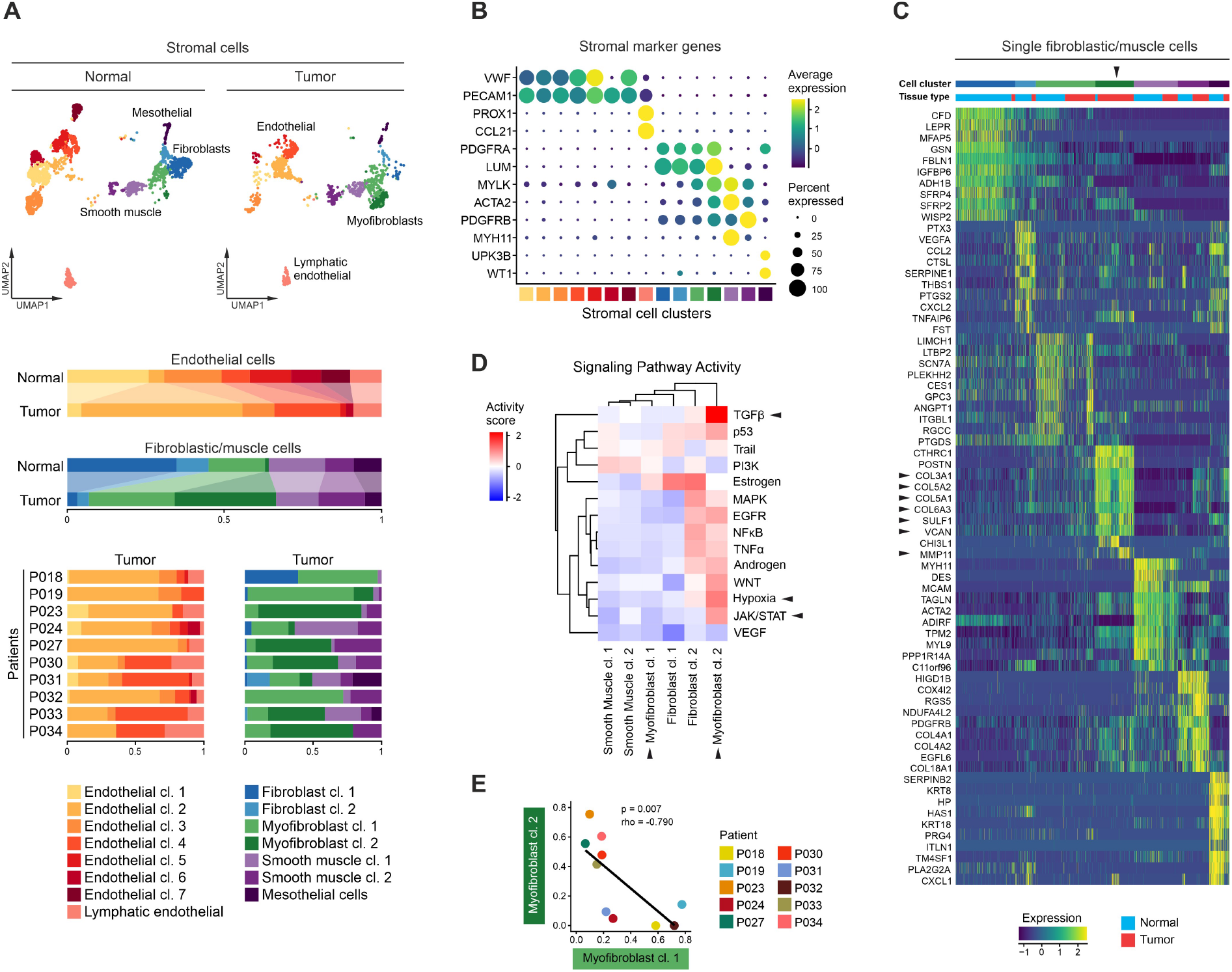
Composition of the stromal microenvironment of lung adenocarcinomas. (A) UMAPs based on the top 20 principal components of all stromal single-cell transcriptomes split by tissue type, color-coded by cell cluster; and relative quantification of endothelial and fibroblastic/muscle cell clusters per tissue type and, for tumor samples, per patient. (B) Average gene expression of selected marker genes for stromal cell clusters, for cell cluster color code see (A). (C) Differentially expressed genes of fibroblastic/muscle cell clusters, maximum top 10 genes showed per cell cluster, for cell cluster color code see (A), black arrowheads indicate relevant marker genes of myofibroblast cluster 2 mentioned in the main text. (D) Mean pathway activity scores of different fibroblastic/muscle cell clusters, mesothelial cells excluded, black arrowheads indicate relevant pathways of myofibroblast clusters 1 and 2 mentioned in the main text. (E) Correlation of the relative quantity of myofibroblast clusters 1 and 2, color-coded by patient; Spearman’s correlation statistics, linear regression line.

Notably, myofibroblast cluster 2 was almost exclusively found in tumor tissues while myofibroblast cluster 1 occurred in both normal and tumor tissues. In myofibroblast cluster 2, differential gene expression analysis revealed high expression of collagens such as *COL3A1*, *COL5A1, COL5A2* and *COL6A3*, other matrix proteins such as *VCAN,* as well as matrix-degrading enzymes such as *SULF1* and *MMP11*, suggesting that these cells are involved in remodeling of the extracellular matrix of the tumor microenvironment (Fig. 3C, arrowheads). Myofibroblast cluster 2 was also characterized by high activity of TGFβ and JAK/STAT signaling as well as hypoxia-induced pathways (Fig. 3D), which are common features of cancer-associated myofibroblasts [18, 19]. In contrast, myofibroblast cluster 1 exhibited low activity scores of all of these pathways. The relative proportion of myofibroblast clusters 1 and 2 within the fibroblastic/muscle cell compartment correlated inversely across patients (Fig. 3E). We conclude that myofibroblasts cluster 1 and 2 represent “normal-like” and “cancer-associated” phenotypes of myofibroblasts, respectively. Moreover, our results suggest tumor subtypes where the tumor stromal microenvironment is predominated by either “normal-like” or “cancer-associated” myofibroblasts.

### The tumor immune microenvironment exhibits pro- and anti-inflammatory traits

We next analyzed immune cells of the tumor microenvironment, as above, to uncover potential roles in both promotion and suppression of tumor growth. We identified different clusters of tissue-resident and monocyte-derived macrophages, monocytes, myeloid and plasmacytoid dendritic cells, mast cells, different clusters of T, NK, B and plasma cells (Fig. 4A, for normal tissue per patient see Supp. Fig. 6A) based on typical marker genes (Fig. 4B) and gene signatures (Supp. Fig. 4A-B) [13, 14]. Although single-cell transcriptomes of most cell clusters were found in both normal and tumor tissue samples, we noted distinct quantitative shifts in the cellular composition of the tumor immune microenvironment.

**Figure 4:**
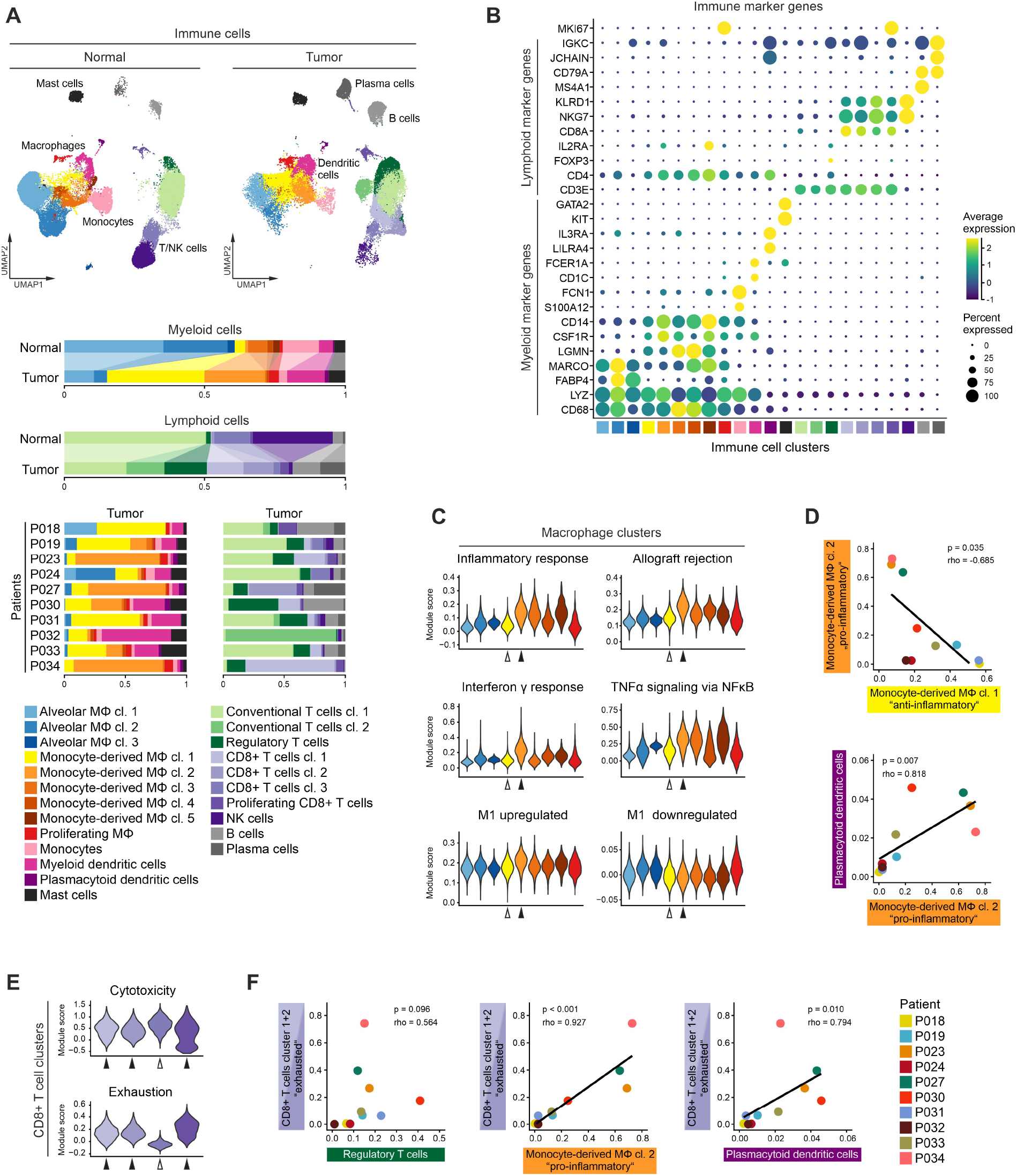
Composition of the immune microenvironment of lung adenocarcinomas. (A) UMAPs based on the top 20 principal components of all immune single-cell transcriptomes split by tissue type, color-coded by cell cluster; and relative quantification of myeloid and lymphoid cell clusters per tissue type and, for tumor samples, per patient. (B) Average gene expression of selected marker genes for immune cell clusters, for cell cluster color code see (A). (C) Module scores of gene signatures related to inflammation and M1/M2 polarization of different macrophage clusters, white and black arrowheads indicate monocyte-derived macrophage clusters 1 and 2, respectively, for cell cluster color code see (A). (D) Correlation of the relative quantity of selected myeloid immune cell clusters, color-coded by patient, for patient color code see (F); Spearman’s correlation statistics, linear regression line. (E) Module scores of gene signatures related to cytotoxicity and exhaustion of different CD8+ T cell clusters, white and black arrowheads indicate cell clusters enriched in normal or tumor tissue, respectively, for cell cluster color code see (A). (F) Correlation of the relative quantity of selected lymphoid and myeloid immune cell clusters, color-coded by patient; Spearman’s correlation statistics, linear regression line.

Among the myeloid cells in tumor tissue samples, we observed an increase in myeloid and plasmacytoid dendritic cells, while monocytes were decreased. Moreover, we found a decrease in tissue-resident and an increase in monocyte-derived macrophages in the tumor samples (Fig. 4A). In addition to conventional macrophage markers, monocyte-derived macrophages were characterized by expression of *CD14, CSF1R* and *LGMN* [20, 21] (Fig. 4B). Among monocyte-derived macrophages, we identified five clusters, of which clusters 1 and 2 clearly predominated the myeloid cell compartment. Here, cluster 2 expressed high levels of proinflammatory chemokines such as *CXCL9* and *CXCL10* and the proinflammatory cytokine *IL1B* (Supp. Fig. 6B, arrowheads). Correspondingly, cluster 2 exhibited high expression scores of gene signatures related to immune response and M1 polarization [22, 23] (Fig. 4C, black arrowheads). In contrast, cluster 1 showed high expression of *SELENOP* (Supp. Fig. 6B, arrowhead), which has been related to M2 polarization of tumor-associated macrophages [24], and low scores of immune response-related signatures (Fig. 4C, white arrowheads). This indicates that monocyte-derived macrophage clusters 1 and 2 represent more anti- and pro-inflammatory functional states, respectively. Notably, the anti- and proinflammatory monocyte-derived macrophage clusters differed between tumor cases (Fig. 4A). The tumor immune microenvironment of some tumors was dominated by the proinflammatory cluster 2 (P023, P027, P030, P034), while in other patients we found predominantly anti-inflammatory cluster 1, or only few monocyte-derived macrophages at all (Fig. 4D). In addition, the proportion of proinflammatory monocyte-derived macrophages cluster 2 correlated with the occurrence of other myeloid cell types such as plasmacytoid dendritic cells (Fig. 4D), pointing at distinct cellular patterns of the myeloid compartment of the tumor immune microenvironment.

Within the lymphoid cell compartment, NK and conventional T cells were decreased, while CD8+ T, B and plasma cells were increased in tumor tissues. In addition, regulatory T cells were almost exclusively found in tumor tissue samples (Fig. 4A). Regulatory T cells expressed inhibitory molecules such as *CTLA4* and *TIGIT* (Supp. Fig. 6C, arrowheads) corresponding to their role in autoimmune tolerance under physiological conditions and immunosuppression in the tumor microenvironment [25]. We identified in total four clusters of CD8+ T cells. Here, we observed a shift from cluster 3 in normal tissues to clusters 1 and 2 and proliferating CD8+ T cells in tumor tissues (Fig. 4A). While all CD8+ T cell clusters showed high expression scores of a gene signature related to cytotoxicity [10], expression scores of an exhaustion signature [10] were only high in the tumor-enriched CD8+ T cell clusters 1 and 2 and proliferating CD8+ T cells (Fig. 4E, black arrowheads), but not in the normal-enriched CD8+ T cell cluster 3 (Fig. 4E, white arrowhead). The abundance of exhausted CD8+ T cells showed a trend towards positive correlation with the proportion of regulatory T cells, while we found a significant positive correlation with proinflammatory monocyte-derived macrophages (cluster 2) and plasmacytoid dendritic cells.

Taken together, we found patient-overarching changes of the tumor immune microenvironment of lung adenocarcinomas such as a shift from tissue-resident to monocyte-derived macrophages and an enrichment in regulatory T cells and exhausted CD8+ T cells. Moreover, our results suggest that interpatient heterogeneity is not restricted to epithelial cancer cells but also reflected by distinct tumor immune microenvironment patterns.

### The tumor microenvironment of lung adenocarcinoma features two major patterns

To integrate our analyses of variable cell prevalences in the tumor microenvironment, we calculated proportions of cells of the myeloid, lymphoid, endothelial and fibroblastic/muscle cell compartments across patients (for cell counts see Supplementary Tables 1-4). Principal component analysis showed that tumors formed subgroups based on the cellular composition of the tumor microenvironment (Fig. 5A). One group of tumors (P018, P019, P024, P031, P032 and P033) was marked by anti-inflammatory monocyte-derived macrophages, NK cells, conventional T cells, myeloid dendritic cells, and normal-like myofibroblasts (referred to as MAN^2^C pattern, Fig. 5B), while a second group of tumors (P023, P027, P030 and P034) was characterized by proinflammatory monocyte-derived macrophages, exhausted CD8+ T cells, plasmacytoid dendritic cells, and cancer-associated myofibroblasts (referred to as CEP^2^ pattern, Fig. 5B). In particular for cell types distinctive of the CEP^2^ pattern, we found many positive correlations of cell type abundances in the tumor microenvironment (Fig. 5C, Supp. Fig. 7A). Furthermore, the two groups based on composition of the microenvironment also largely separated histologically well and moderately differentiated tumors from more poorly differentiated tumors (Fig. 5A). This was also reflected by mean expression scores of the “alveolar/club-like” or “undifferentiated” tumor cell signatures, respectively (Fig. 5B). These results illustrate that lung adenocarcinomas harbor two distinct tumor microenvironment patterns, termed MAN^2^C and CEP^2^, correlating with high and low tumor differentiation grade, respectively.

**Figure 5:**
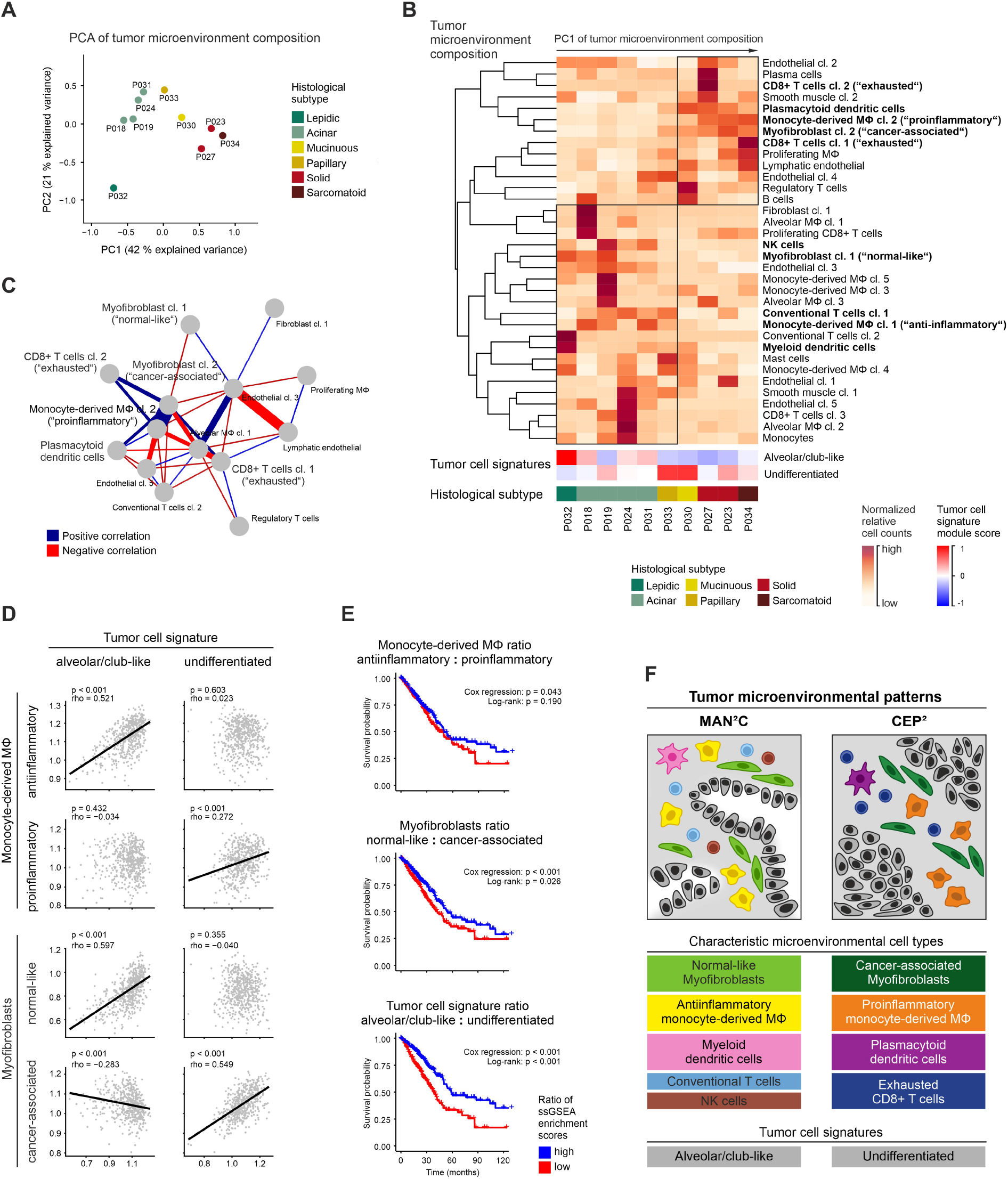
Tumor microenvironmental patterns in lung adenocarcinomas. (A) Principal component analysis based on the proportion of stromal and immune cell clusters, color-coded by histological subtype, patients indicated. (B) Normalized proportion of stromal and immune cell clusters, mean module scores of tumor cell signature, and histological subtypes per patient, patients sorted along the first principal component from principal component analysis in (A), cell clusters included in the model in (F) in bold. (C) Correlation of the proportion of stromal and immune cell clusters, most connected section of correlation network plot shown; Spearman’s correlation statistics, only correlations with rho > 0.7 and p < 0.05 shown. (A+B+C) Cell clusters occurring in less than 3 patients were excluded from analyses. (D) Correlation of single-sample gene set enrichment analysis (ssGSEA) enrichment scores of 533 patients from the TCGA lung adenocarcinoma cohort based on marker genes of selected immune and stromal cell subtypes and tumor cell signatures; Spearman’s correlation statistics, linear regression line. (E) Kaplan-Meier overall survival curves of 524 patients from the TCGA lung adenocarcinoma cohort grouped by ratios of ssGSEA enrichment scores for selected immune and stromal cell subtypes or tumor cell signatures; log-rank statistics, univariate Cox regression. (F) Schematic representation of subtypes of lung adenocarcinoma characterized by different grades of tumor epithelial cell differentiation and different composition of the corresponding tumor microenvironment.

To test if these tumor microenvironmental expression patterns can be recapitulated in larger patient cohorts, we analyzed bulk gene expression data of The Cancer Genome Atlas (TCGA) lung adenocarcinoma cohort. Expression of cell type signatures identified by single-cell RNA sequencing was quantified in the bulk data by single-sample gene set enrichment analysis (ssGSEA) [26] (for marker genes see Supplementary Tables 5-8). We found a specific positive correlation of the alveolar/club-like tumor cell signature with anti-inflammatory monocyte-derived macrophages and normal-like myofibroblasts, recapitulating the MAN^2^C pattern, and, on the other hand, a positive correlation of the undifferentiated tumor cell signature with proinflammatory monocyte-derived macrophages and cancer-associated myofibroblasts, recapitulating the CEP^2^ pattern (Fig. 5D, Supp. Fig. 7B). Hence, we were able to recapitulate in a large cohort the distribution of macrophage and myofibroblast subtypes of the tumor microenvironment patterns revealed by single-cell RNA sequencing, although analysis of bulk data may miss scarce cell types and is likely to be influenced by relative sizes of the immune or stromal compartments in the bulk samples.

To investigate the biological and clinical relevance of these patient subgroups, we analyzed the overall survival of the TCGA lung adenocarcinoma cohort contigent on the different gene signatures of tumor epithelial cells, monocyte-derived macrophages and myofibroblasts. Since our single-cell analyses revealed that alveolar/club-like and undifferentiated tumor cells, anti- and pro-inflammatory macrophages, and normal-like and cancer-associated myofibroblasts were inversely correlated, respectively, we used the ratio of corresponding signatures to stratify patients. Gene signatures representing the MAN^2^C pattern, that is alveolar/club-like tumor cells, anti-inflammatory monocyte-derived macrophages and normal-like myofibroblasts, were associated with a better overall survival compared to the CEP^2^ pattern, represented by gene signatures of undifferentiated tumor cells, pro-inflammatory monocyte-derived macrophages and cancer-associated myofibroblasts (Fig. 5E). This association was confirmed in univariate analyses for all signatures (tumor cell signatures: HR 0.10, 95%CI 0.03-0.26, p < 0.001, myofibroblast signatures: HR 0.08, 95%CI 0.02-0.28, p < 0.001, macrophage signatures: HR 0.14, 95%CI 0.02-0.94, p = 0.043) and in multivariate analyses for the tumor cell signatures (HR 0.13, 95%CI 0.03-0.56, p < 0.001), hinting at a potential clinical relevance of the MAN^2^C and CEP^2^ patterns in lung adenocarcinoma.

In summary, our results show that lung adenocarcinomas can be stratified into clinically relevant subgroups which are not only characterized by the grade of tumor epithelial cells, but also by the cellular composition of their associated tumor microenvironment (Fig. 5F). We therefore propose two major microenvironmental patterns in lung adenocarcinoma which we term MAN^2^C and CEP^2^ according to the initial letters of the respective characteristic cell types.

## Discussion

By applying single-cell RNA sequencing to lung adenocarcinomas, we identified different cell types enriched in the tumor microenvironment such as myofibroblasts, monocyte-derived macrophages, dendritic cells, regulatory T cells and exhausted CD8+ T cells, and observed a high interpatient heterogeneity with respect to their distribution. This led us to identify two major microenvironmental patterns: i) a rather normal-like tumor microenvironment characterized by normal-like myofibroblasts, conventional T cells, NK cells, but also anti-inflammatory monocyte-derived macrophages and myeloid dendritic cells, referred to as MAN^2^C pattern, and ii) a more distinctly altered tumor microenvironment characterized by proinflammatory monocyte-derived macrophages, plasmacytoid dendritic cells, exhausted CD8+ T cells and cancer-associated myofibroblasts, referred to as CEP^2^ pattern (Fig. 5F). Here, the MAN^2^C pattern was associated with an alveolar/club-like gene expression pattern of carcinoma cells, lower histological grade and better prognosis, while the CEP^2^ pattern was associated with an undifferentiated gene expression pattern of carcinoma cells, higher histological grade and worse prognosis.

Currently, besides tumor stage, histological grading based solely on the predominant tumor growth pattern is used to predict prognosis in lung adenocarcinoma. However, in particular for intermediate grade tumors, a more precise risk stratification is needed [27]. Different histological features, including nuclear grade, mitotic count, and tumor budding [27], as well as a combined histological grading taking into account intertumoral heterogeneity [28] have been proposed. However, these concepts focus on the carcinoma cells and do not include potentially prognostically relevant features of the complex composition of the tumor microenvironment. Most available data focus on single cell types such as tumor-associated macrophages. Although evidence points to a positive prognostic effect of pro-inflammatory tumor-associated macrophages (M1 macrophages), results from many studies are inconclusive which might be due to the fact that differentiation between macrophage subtypes based on a limited number of immunohistochemical markers is not straightforward [29]. Our results suggest that the microenvironmental CEP^2^ pattern is associated with worse prognosis. While this contradicts some previous works with regard to pro-inflammatory tumor-associated macrophages [29], it is in line with existing evidence for cancer-associated myofibroblasts [30] and plasmacytoid dendritic cells [31]. The absence of cancer-associated myofibroblasts, and the presence of NK cells [32] and conventional T cells [33] support the association of the MAN^2^C pattern with better prognosis. In contrast to recent evaluations of the tumor microenvironment by bulk gene expression profiling [34–36], our analysis proposes novel microenvironmental signatures by comprehensive in-depth characterization of different immune and stromal cells complementing the carcinoma cell-centric view.

As immune cells have already been shown to have predictive value underlined by the introduction of immune checkpoint inhibitors and the evaluation of PD-L1 expression in some cancers (although not yet lung cancer [37]), it is likely that more comprehensive characterizations of the immune tumor microenvironment will provide even deeper insights in patient stratification and drug development. Recently, expression of the proinflammatory cytokines CXCL9 and CXCL10 by tumor-associated macrophages, for instance, has been shown to be essential for tumor response to anti-PD-L1 therapy [38, 39]. Interestingly, we observed high expression of these cytokines in pro-inflammatory monocyte-derived macrophages indicating that the microenvironmental CEP^2^ pattern, which was also marked by exhausted CD8+ T cells, could be a predictor of response to anti-PD-L1 therapy. In addition, the CEP^2^ pattern was characterized by cancer-associated myofibroblasts which showed high expression of potential novel therapeutic targets, such as *PSTN* and *MMP11* [40, 41]. Despite an overall depletion of NK cells in tumors in agreement with other studies [7, 9], the MAN^2^C microenvironmental pattern still contained small populations of NK cells which could be targeted by immunostimulatory agents [42]. Although not specifically related to the MAN^2^C or CEP^2^ pattern, we and others [8, 9] observed an enrichment of B cells in some tumors with potential importance in development of novel immunotherapies [43].

While the analysis of the tumor microenvironment will likely benefit from single cell approaches most substantially because of its complex cellular composition, they will also contribute to a better understanding of the tumor itself because lung cancers are characterized by high tumor mutational burden [44] and high genomic intertumoral heterogeneity [45]. On the transcriptional level, differences have been described between squamous and adenocarcinomas of the lung [46], while this is less evident for histological subtypes of lung adenocarcinoma [47]. Interestingly, despite intertumoral heterogeneity, we found that single-cell transcriptomes of tumor epithelial cells retained similarities across patients corresponding to the established histological subtypes of lung adenocarcinoma [27]. We defined two inversely correlated tumor cell signatures, “alveolar/club-like” and “undifferentiated”, reflecting a transcriptional differentiation gradient reminiscent of transcriptional patterns, recently described by Kim and colleagues [9]. Furthermore, the “undifferentiated” tumor cell signature was associated with high EGFR signaling, being already a therapeutic target in *KRAS*-wildtype lung adenocarcinomas [4], and JAK/STAT signaling, which is a possible therapeutic target in preclinical *KRAS*-mutant tumor models [48]. The “alveolar/club-like” tumor cell signature correlated with high PI3K pathway activity, which has been under early clinical investigation as therapeutic target, however, with yet unconvincing results, possibly due to insufficient molecular characterization and selection of patients [49].

Our findings show that single cell approaches contribute to a better understanding of the heterogeneity within each tumor tissue compartment, but may also prove invaluable for analysing the complex interactions between malignant epithelial cells and various non-malignant stromal and immune cells. In our study, we identified two major tumor microenvironmental patterns associated with tumor differentiation grade, termed MAN^2^C and CEP^2^, with the potential to predict prognosis, therapy response and provide insight on possible novel therapeutic targets. Nevertheless, the relevance of our proposed tumor microenvironmental subtypes will have to be validated in larger clinical cohorts. Moreover, the question of cause and consequence remains open: To which extent do cancer cells shape their microenvironment or does the microenvironment affect cancer cells? Given the cellular diversity of the tumor microenvironment, many possible cellular interactions have been studied in preclinical models. These preclinical efforts need to be complemented by translational studies to identify critical mechanisms in this complex network that determine tumor response to targeted or immune therapies in the clinical context. While our study demonstrates how single-cell gene expression profiling of clinical samples can contribute to this task, in the future, other single-cell approaches comprising spatial information [50], surface protein expression [51] and epigenetic characterization [52] will help to complete the picture [53].

## Methods

### Collection of tissue specimens

Fresh normal lung parenchyma and tumor tissues were obtained from previously untreated lung adenocarcinoma patients undergoing primary surgery during intraoperative pathologist consultation. All patients were aware of the planned research and agreed to the use of tissue. Research was approved by vote EA4/164/19 of the ethics committee of Charité - Universitätsmedizin Berlin.

### Tissue dissociation and single cell isolation

Tissue specimens of approximately 0.1-0.5 cm^3^ were stored short-term (maximum 3 hours) on ice in Tissue Storage Solution (Miltenyi, 130-100-008) for transport. Next, tissues were minced using two scalpels and dissociated using the Tumor Dissociation Kit, human (Miltenyi, 130-095-929) and a gentleMACS Octo Dissociator with heaters (Miltenyi, 130-096-427), using program 37C_h_TDK_1 for 30-45 min. Cell suspensions were filtered using 100 μm filters, and all subsequent steps were performed at 4°C or on ice. Cells were pelleted by centrifugation at 300 g for 5 min in BSA-coated low-binding tubes, and cells were treated with 1 ml ACK erythrocyte lysis buffer for 60 seconds and washed with DMEM. Cells were pelleted, resuspended in PBS, cell suspensions were filtered using 20 μm filters, debris was removed using the Debris Removal Solution (Miltenyi, 130-109-398), and cells were counted using a Neubauer chamber.

### Single-cell RNA sequencing

10,000 single cells were used for single-cell library production, using the Chromium Single Cell 3’Reagent Kit v3 and the Chromium Controller (10x Genomics) according to the manufacturer’s protocol. Libraries were sequenced on a HiSeq 4000 Sequencer (Illumina) at 100-400 mio. reads per library to a mean library saturation of 55% resulting in average 54,000 reads per cell.

### H&E and immunostaining

3-5 μm tissue sections formalin-fixed and paraffin-embedded (FFPE) tissue were used for immunofluorescence, immunohistochemistry and hematoxylin and eosin (H&E) staining. For H&E staining, sections were stained for 8 min in acidic haemalum staining solution (Waldeck) and for 2,5 min in eosin staining solution (Sigma-Aldrich) at room temperature using a Tissue-Tek Prisma Plus slide stainer (Sakura).

Immunohistochemical and immunofluorescence stainings of FFPE tissue sections were performed on the BenchMark XT immunostainer (Ventana Medical Systems). For antigen retrieval, tissue sections were incubated in CC1 or CC2 buffer (both Ventana Medical Systems) for 30 min at 100°C. Sections were incubated with primary antibodies for 60 min and with secondary antibodies for 30 minutes at room temperature diluted in Dako Real Antibody Diluent (Dako, S2022). The following primary antibodies were used: rabbit anti-TFF3 (Abcam, ab108599, 1:250, CC1 antigen retrieval), rabbit anti-EGFR (Roche, #790-4347, prediluted, CC1 antigen retrieval), mouse anti-p16 (Roche, #805-4713, prediluted, CC1 antigen retrieval), mouse anti-SFTPA (Abcam, ab51891, 1:200, CC2 antigen retrieval). Hematoxylin-and-eosin and immunohistochemical images were taken using a Pannoramic SCAN 150 slide scanner (3DHISTECH).

### Panel sequencing

For panel sequencing of oncogenic mutations and gene fusions, tumor-enriched areas were macrodissected from formalin-fixed and paraffin-embedded tissue sections.

DNA was isolated using the Maxwell RSC DNA FFPE Kit (Promega) on a Maxwell RSC 48 Instrument (Promega). Isolated DNA was analyzed for oncogenic mutations using the nNGM panel v1 (ThermoFisher), an Ion 530 chip (ThermoFisher) and the Ion Chef/Ion S5 XL System (ThermoFisher).

RNA was isolated using the Maxwell RSC RNA FFPE Kit (Promega) on a Maxwell RSC 48 Instrument (Promega). Isolated RNA was analyzed for oncogenic gene fusions using the Oncomine Focus RNA Assay (ThermoFisher), an Ion 530 chip and the Ion Chef/Ion S5 XL System (ThermoFisher).

The Sequence Pilot Software (Version 4.4.0, JSI Medical Systems) and the Ion Reporter Software (Version 5.12, ThermoFisher) were used for variant calling.

### Single-cell RNA sequencing data analysis

#### Preprocessing, filtering and normalization

For each sample, UMIs were quantified using Cellranger 3.0.2 (10x Genomics) with reference transcriptome GRCh38. Subsequent analyses were performed using the R package “Seurat v3” [54], if not stated otherwise. Single-cell gene expression data of all patients were merged. Quality control filters were set to only include cells with 500-10,000 genes detected, 1,000-100,000 UMIs counted, fraction of mitochondrial reads <30%, and fraction of hemoglobin reads <5%. After filtering, UMI counts were variance-stabilized using scTransform with 3,000 variable features [55], while regressing out number of UMIs counted and fraction of mitochondrial reads. Clustering and cell type annotation

The top 15 principal components were used to construct shared nearest neighbor (SNN) graph and UMAP embedding as implemented in the R package “Seurat v3” with default parameters. Next, main cell types (epithelial, immune, stromal) were identified by scoring cell type markers across clusters (resolution = 0.2, otherwise default parameters). PCA, SNN graph construction and UMAP embedding was rerun on each subset using the top 20 principal components, and a resolution of 1 for the epithelial and stromal subsets, or 0.5 for the immune subset with otherwise default parameters. Cell type markers used to score epithelial, immune, and stromal cell types were adapted from Habermann et al. [56] and Tata et al. [57]. Cell type signatures from Vieira Braga et al. [13] and Travaglini et al. [14] were used to validate manual cell type annotation. Epithelial or immune contaminated clusters were identified by expression of *EPCAM* or *PTPRC*, respectively, and removed prior to further analyses. In the epithelial subset, cell clusters which were relatively overrepresented in the tumor tissue samples were annotated as tumor cells.

#### Differential gene expression analysis

For differential gene expression analysis of epithelial cells, tumor cells from tumor samples were subset and gene expression rescaled. Immune and stromal subsets were split as stated, and gene expression rescaled. Marker genes for each cell cluster versus all cells of the subset were computed using the FindAllMarkers function of the R package “Seurat v3” with Wilcoxon rank-sum test and the following parameters: only positive markers, fraction of expressing cells inside the cluster >0.25, difference between fraction of expressing cells inside and outside the cluster >0.25, log fold change between cells inside and outside the cluster >0.25.

#### Functional analysis

Cell cycle phases were scores as implemented in the R package “Seurat v3”. Expression of gene sets of the Hallmark signature collection of the Broad Institute [22], and M1 vs. M2 up- and downregulated genes [23] were scored using the AddModuleScore function of the R package “Seurat v3”. Oncogenic signaling pathway activity scores were computed using the R package “progeny” [15, 16].

### TCGA gene expression analysis

FPKM-normalized gene expression values from RNA sequencing and clinical data of the lung adenocarcinoma (LUAD) cohort of The Cancer Genome Project was downloaded using the R package “TCGAbiolinks” [58] and log2 transformed.

Data from single-cell RNA sequencing was subset into myeloid, lymphoid, endothelial and stromal compartments, rescaled using the ScaleData function and marker genes identified using the FindAllMarkers function (only positive genes, fraction of expressing cells ≥ 0.25, difference of fraction of expressing cells ≥ 0.25).

For epithelial marker genes, tumor-specific clusters in tumor samples were subset, rescaled using the ScaleData function and dimensionality reduction performed using RunPCA. The top 30 genes positively or negatively correlated with principal component 1 were selected for the PC1^pos^ or PC1^neg^ marker gene list, respectively.

Marker gene lists from myeloid, lymphoid, endothelial, stromal and epithelial cell subsets were used as gene sets to perform single-sample gene set enrichment analysis (ssGSEA) [26] on TCGA LUAD gene expression data using the R package “GSVA” [59] assuming Gaussian distribution.

Associations between enrichment scores (ES) from ssGSEA were calculated by Pearson correlation test using the R package “corrplot”.

### TCGA survival analysis

For survival analyses, ES from ssGSEA of the TCGA LUAD gene expression data were dichotomized (ES > median or ≤ median). For dichotomization of combined enrichment scores, the enrichment score matrix was rotated clockwise by the arctangent of the slope from linear regression using a rotation matrix and subsequently dichotomized (x value > median or ≤ median).

For survival analysis, survival curves, log rank statistics and Cox regression were calculated including the parameters “vital status”, “days to death” and “days to last follow up” from the TCGA dataset using the R packages “survival” and “survminer”.

## Author contributions

NB, FK, PB conceived and designed the study;

PB, AT, FK, DH, AE, TD, JN, JR contributed to clinical sample acquisition and preparation;

AT, PB conducted experiments;

PB, BO, JP, DB, NB performed bioinformatic analyses;

PB, FK, NB, PJ, CF, DH, MM, CS analyzed and interpreted data and/or supervised parts of the study;

PB wrote the manuscript;

FK, NB revised the manuscript;

all authors provided critical feedback and helped shaping the research, analysis, and manuscript.

## Acknowledgement

We thank Manuela Pacyna-Gengelbach and Barabara Meyer-Bartell for excellent technical assistance.

The results of this study are in part based upon data generated by the TCGA Research Network: https://www.cancer.gov/tcga.

## References

1. Bray, F., et al., Global cancer statistics 2018: GLOBOCAN estimates of incidence and mortality worldwide for 36 cancers in 185 countries. CA Cancer J Clin, 2018. 68(6): p. 394–424.

2. Wakelee, H.A., et al., Lung cancer incidence in never smokers. J Clin Oncol, 2007. 25(5): p. 472–8.

3. Cancer Genome Atlas Research, N., Comprehensive molecular profiling of lung adenocarcinoma. Nature, 2014. 511(7511): p. 543–50.

4. Hirsch, F.R., et al., Lung cancer: current therapies and new targeted treatments. Lancet, 2017. 389(10066): p. 299–311.

5. Memon, H. and B.M. Patel, Immune checkpoint inhibitors in non-small cell lung cancer: A bird’s eye view. Life Sci, 2019. 233: p. 116713.

6. Altorki, N.K., et al., The lung microenvironment: an important regulator of tumour growth and metastasis. Nat Rev Cancer, 2019. 19(1): p. 9–31.

7. Lavin, Y., et al., Innate Immune Landscape in Early Lung Adenocarcinoma by Paired Single-Cell Analyses. Cell, 2017. 169(4): p. 750–765 e17.

8. Lambrechts, D., et al., Phenotype molding of stromal cells in the lung tumor microenvironment. Nat Med, 2018. 24(8): p. 1277–1289.

9. Kim, N., et al., Single-cell RNA sequencing demonstrates the molecular and cellular reprogramming of metastatic lung adenocarcinoma. Nat Commun, 2020. 11(1): p. 2285.

10. Guo, X., et al., Global characterization of T cells in non-small-cell lung cancer by single-cell sequencing. Nat Med, 2018. 24(7): p. 978–985.

11. Becht, E., et al., Dimensionality reduction for visualizing single-cell data using UMAP. Nat Biotechnol, 2018.

12. Lee, H.O., et al., Lineage-dependent gene expression programs influence the immune landscape of colorectal cancer. Nat Genet, 2020.

13. Vieira Braga, F.A., et al., A cellular census of human lungs identifies novel cell states in health and in asthma. Nat Med, 2019. 25(7): p. 1153–1163.

14. Travaglini, K.J., et al., A molecular cell atlas of the human lung from single-cell RNA sequencing. Nature, 2020.

15. Schubert, M., et al., Perturbation-response genes reveal signaling footprints in cancer gene expression. Nat Commun, 2018. 9(1): p. 20.

16. Holland, C.H., et al., Robustness and applicability of transcription factor and pathway analysis tools on single-cell RNA-seq data. Genome Biol, 2020. 21(1): p. 36.

17. Naizhen, X., et al., Spatiotemporal Expression of Three Secretoglobin Proteins, SCGB1A1, SCGB3A1, and SCGB3A2, in Mouse Airway Epithelia. J Histochem Cytochem, 2019. 67(6): p. 453–463.

18. Yoshida, G.J., Regulation of heterogeneous cancer-associated fibroblasts: the molecular pathology of activated signaling pathways. J Exp Clin Cancer Res, 2020. 39(1): p. 112.

19. Petrova, V., et al., The hypoxic tumour microenvironment. Oncogenesis, 2018. 7(1): p. 10.

20. Franklin, R.A. and M.O. Li, Ontogeny of Tumor-associated Macrophages and Its Implication in Cancer Regulation. Trends Cancer, 2016. 2(1): p. 20–34.

21. Solberg, R., et al., Legumain expression, activity and secretion are increased during monocyte-to-macrophage differentiation and inhibited by atorvastatin. Biol Chem, 2015. 396(1): p. 71–80.

22. Liberzon, A., et al., The Molecular Signatures Database (MSigDB) hallmark gene set collection. Cell Syst, 2015. 1(6): p. 417–425.

23. Martinez, F.O., et al., Transcriptional profiling of the human monocyte-to-macrophage differentiation and polarization: new molecules and patterns of gene expression. J Immunol, 2006. 177(10): p. 7303–11.

24. Solinas, G., et al., Tumor-conditioned macrophages secrete migration-stimulating factor: a new marker for M2-polarization, influencing tumor cell motility. J Immunol, 2010. 185(1): p. 642–52.

25. Jiang, Y., Y. Li, and B. Zhu, T-cell exhaustion in the tumor microenvironment. Cell Death Dis, 2015. 6: p. e1792.

26. Barbie, D.A., et al., Systematic RNA interference reveals that oncogenic KRAS-driven cancers require TBK1. Nature, 2009. 462(7269): p. 108–12.

27. Travis, W.D., et al., The 2015 World Health Organization Classification of Lung Tumors: Impact of Genetic, Clinical and Radiologic Advances Since the 2004 Classification. J Thorac Oncol, 2015. 10(9): p. 1243–1260.

28. Moreira, A.L., et al., A Grading System for Invasive Pulmonary Adenocarcinoma: A Proposal From the International Association for the Study of Lung Cancer Pathology Committee. J Thorac Oncol, 2020. 15(10): p. 1599–1610.

29. Conway, E.M., et al., Macrophages, Inflammation, and Lung Cancer. Am J Respir Crit Care Med, 2016. 193(2): p. 116–30.

30. Ito, M., et al., Prognostic impact of cancer-associated stromal cells in patients with stage I lung adenocarcinoma. Chest, 2012. 142(1): p. 151–158.

31. Koucky, V., J. Boucek, and A. Fialova, Immunology of Plasmacytoid Dendritic Cells in Solid Tumors: A Brief Review. Cancers (Basel), 2019. 11(4).

32. Zhang, S., et al., Prognostic Significance of Tumor-Infiltrating Natural Killer Cells in Solid Tumors: A Systematic Review and Meta-Analysis. Front Immunol, 2020. 11: p. 1242.

33. Geng, Y., et al., Prognostic Role of Tumor-Infiltrating Lymphocytes in Lung Cancer: a Meta-Analysis. Cell Physiol Biochem, 2015. 37(4): p. 1560–71.

34. Tao, Y., Y. Li, and B. Liang, Comprehensive analysis of microenvironment-related genes in lung adenocarcinoma. Future Oncol, 2020. 16(24): p. 1825–1837.

35. Ojlert, A.K., et al., The immune microenvironment in non-small cell lung cancer is predictive of prognosis after surgery. Mol Oncol, 2019. 13(5): p. 1166–1179.

36. Huang, J., et al., Tumor microenvironment characterization identifies two lung adenocarcinoma subtypes with specific immune and metabolic state. Cancer Sci, 2020. 111(6): p. 1876–1886.

37. Bironzo, P. and M. Di Maio, A review of guidelines for lung cancer. J Thorac Dis, 2018. 10(Suppl 13): p. S1556–S1563.

38. Qu, Y., et al., Baseline Frequency of Inflammatory Cxcl9-Expressing Tumor-Associated Macrophages Predicts Response to Avelumab Treatment. Cell Rep, 2020. 32(1): p. 107873.

39. House, I.G., et al., Macrophage-Derived CXCL9 and CXCL10 Are Required for Antitumor Immune Responses Following Immune Checkpoint Blockade. Clin Cancer Res, 2020. 26(2): p. 487–504.

40. Okazaki, T., et al., Periostin is a negative prognostic factor and promotes cancer cell proliferation in non-small cell lung cancer. Oncotarget, 2018. 9(58): p. 31187–31199.

41. Yang, H., et al., Matrix Metalloproteinase 11 Is a Potential Therapeutic Target in Lung Adenocarcinoma. Mol Ther Oncolytics, 2019. 14: p. 82–93.

42. Wrangle, J.M., et al., ALT-803, an IL-15 superagonist, in combination with nivolumab in patients with metastatic non-small cell lung cancer: a non-randomised, open-label, phase 1b trial. Lancet Oncol, 2018. 19(5): p. 694–704.

43. Patel, A.J., et al., The role of B lymphocytes in the immuno-biology of non-small-cell lung cancer. Cancer Immunol Immunother, 2020. 69(3): p. 325–342.

44. Alexandrov, L.B., et al., Signatures of mutational processes in human cancer. Nature, 2013. 500(7463): p. 415–21.

45. Liu, Y., et al., Genomic heterogeneity of multiple synchronous lung cancer. Nat Commun, 2016. 7: p. 13200.

46. Niemira, M., et al., Molecular Signature of Subtypes of Non-Small-Cell Lung Cancer by Large-Scale Transcriptional Profiling: Identification of Key Modules and Genes by Weighted Gene Co-Expression Network Analysis (WGCNA). Cancers (Basel), 2019. 12(1).

47. Molina-Romero, C., et al., Differential gene expression profiles according to the Association for the Study of Lung Cancer/American Thoracic Society/European Respiratory Society histopathological classification in lung adenocarcinoma subtypes. Hum Pathol, 2017. 66: p. 188–199.

48. Mohrherr, J., et al., JAK-STAT inhibition impairs K-RAS-driven lung adenocarcinoma progression. Int J Cancer, 2019. 145(12): p. 3376–3388.

49. Tan, A.C., Targeting the PI3K/Akt/mTOR pathway in non-small cell lung cancer (NSCLC). Thorac Cancer, 2020. 11(3): p. 511–518.

50. Stahl, P.L., et al., Visualization and analysis of gene expression in tissue sections by spatial transcriptomics. Science, 2016. 353(6294): p. 78–82.

51. Stoeckius, M., et al., Simultaneous epitope and transcriptome measurement in single cells. Nat Methods, 2017. 14(9): p. 865–868.

52. Buenrostro, J.D., et al., Single-cell chromatin accessibility reveals principles of regulatory variation. Nature, 2015. 523(7561): p. 486–90.

53. Rajewsky, N., et al., LifeTime and improving European healthcare through cell-based interceptive medicine. Nature, 2020.

54. Stuart, T., et al., Comprehensive Integration of Single-Cell Data. Cell, 2019. 177(7): p. 1888–1902 e21.

55. Hafemeister, C. and R. Satija, Normalization and variance stabilization of single-cell RNA-seq data using regularized negative binomial regression. Genome Biol, 2019. 20(1): p. 296.

56. Habermann, A.C., et al., Single-cell RNA sequencing reveals profibrotic roles of distinct epithelial and mesenchymal lineages in pulmonary fibrosis. Sci Adv, 2020. 6(28): p. eaba1972.

57. Tata, P.R. and J. Rajagopal, Plasticity in the lung: making and breaking cell identity. Development, 2017. 144(5): p. 755–766.

58. Colaprico, A., et al., TCGAbiolinks: an R/Bioconductor package for integrative analysis of TCGA data. Nucleic Acids Res, 2016. 44(8): p. e71.

59. Hanzelmann, S., R. Castelo, and J. Guinney, GSVA: gene set variation analysis for microarray and RNA-seq data. BMC Bioinformatics, 2013. 14: p. 7.

