## Supplemental Figures 1-7 for "Single-cell RNA sequencing reveals distinct tumor microenvironmental patterns in lung adenocarcinoma"

[#] joint last authors

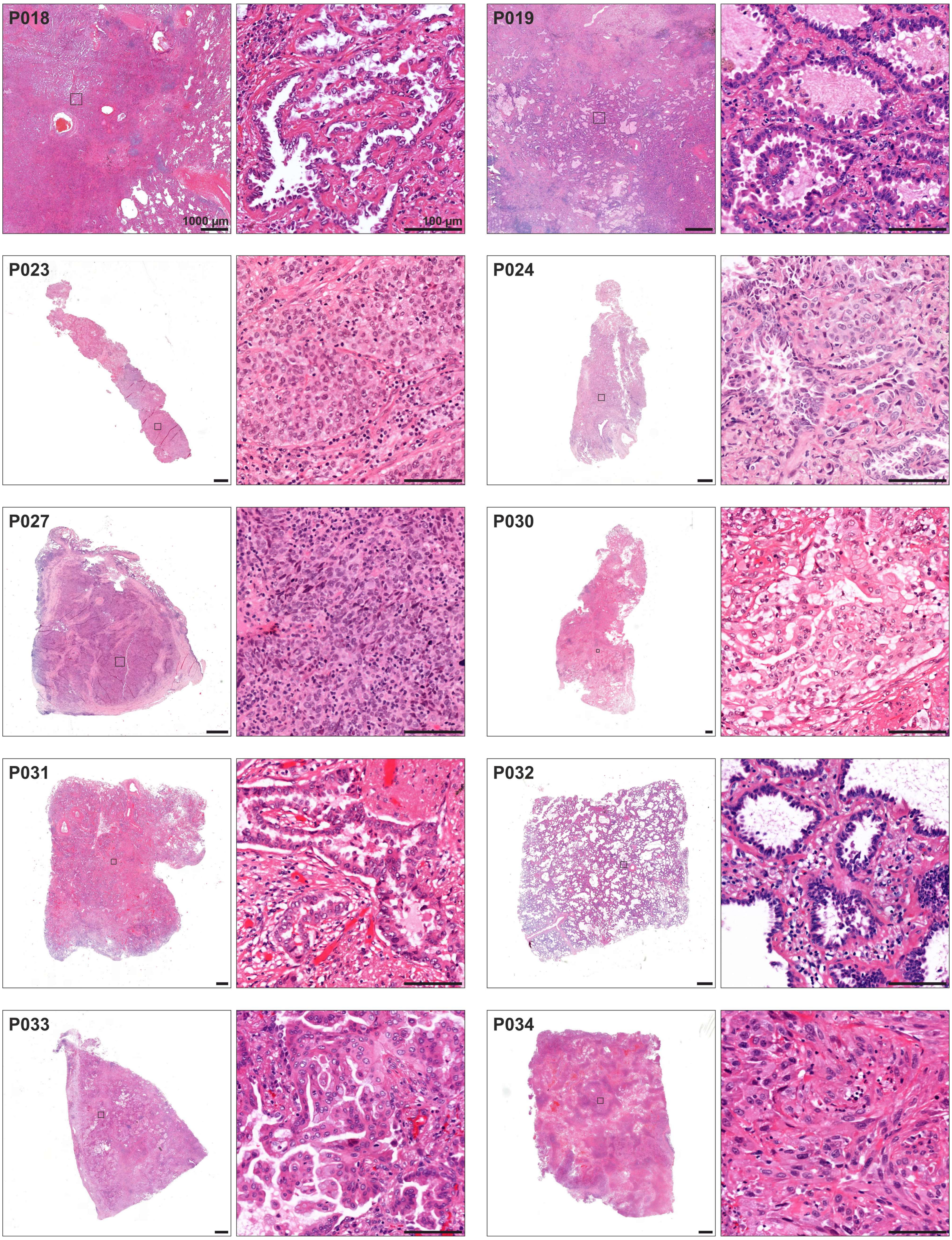


**Supplemental Figure 1**

H&E staining of lung adenocarcinomas analyzed in the study. For patients P018 and P019, representative tumor tissue shown, for all other patients, consecutive tissue sections of the sample used to prepare single-cell suspensions is shown. Black squares indicate magnified areas. Scale bars indicate 1000 µm or 100 µm in overview or magnified images, respectively.

A

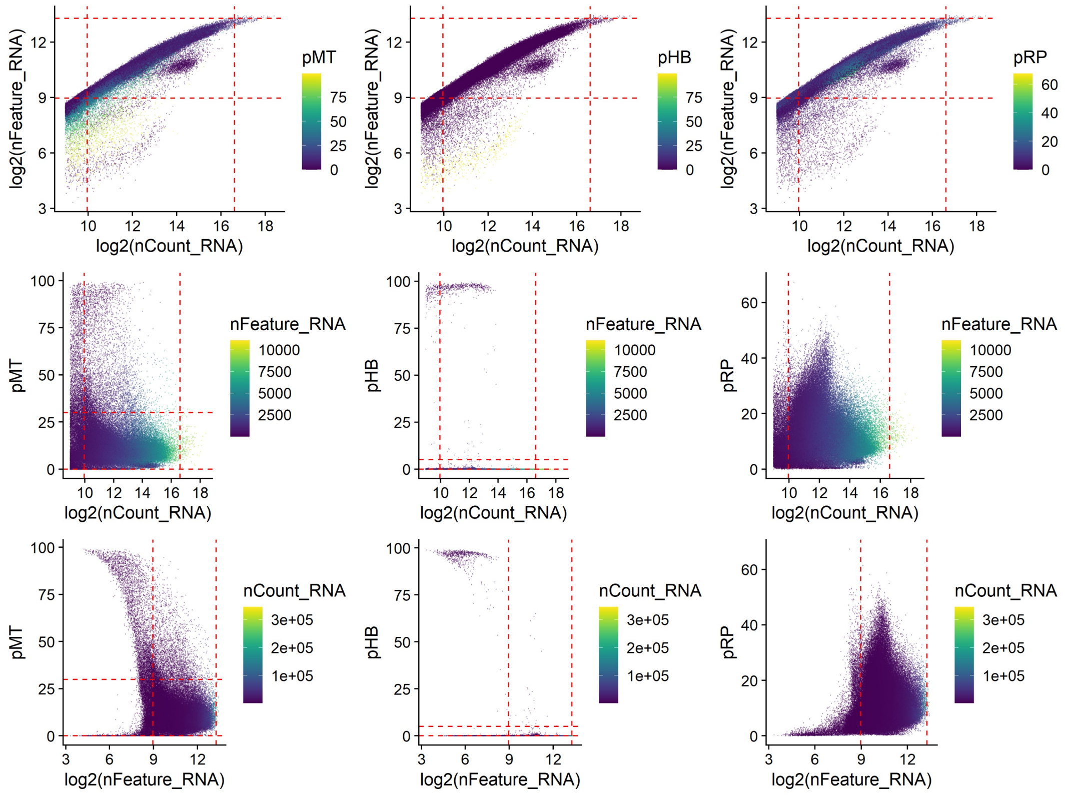


B

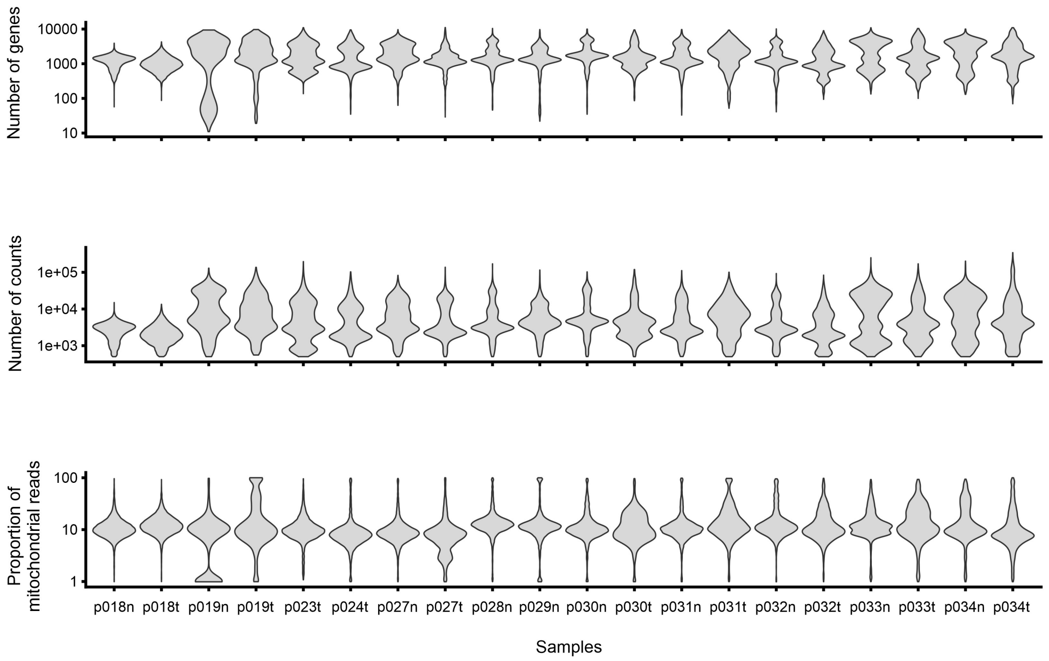


**Supplemental Figure 2**

(A) Scatter plots of quality control parameters of all single-cell transcriptomes analyzed in the study, red dashed lines indicate cut-off values for filtering of high-quality transcriptomes. (B) Violin plots of selected quality control parameters grouped by tissue sample, n = normal tissue, t = tumor tissue.

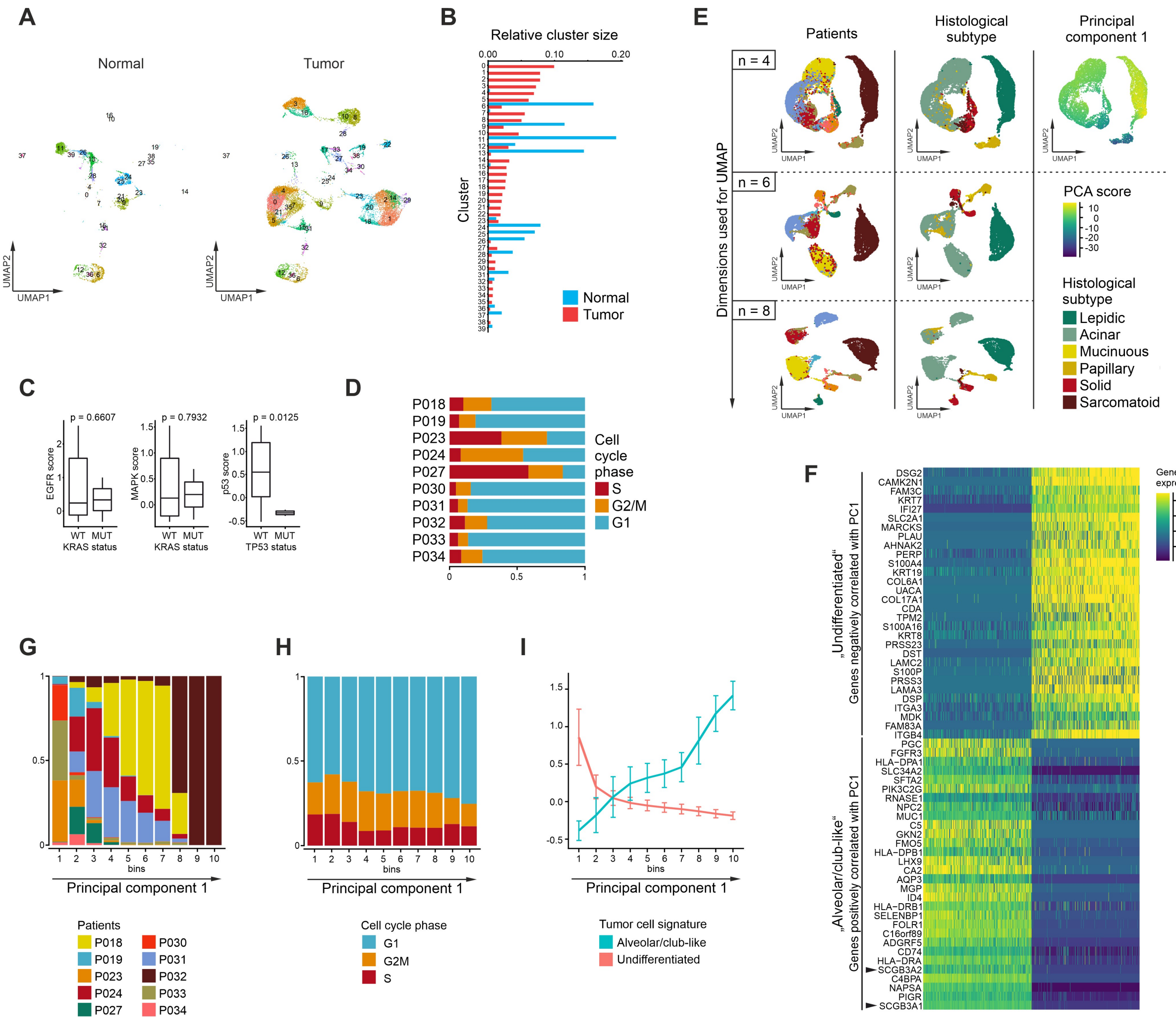


**Supplemental Figure 3**

(A) UMAPs based on the top 20 principal components of all epithelial single-cell transcriptomes split by tissue type, color-coded by cluster, cluster number indicated. (B) Relative size of clusters within normal (blue) or tumor (red) tissue samples, respectively. (C) Pathway activity scores of selected signaling pathways in KRAS wildtype and mutated, or TP53 wildtype and mutated patients, respectively; Welch's t-test statistics. (D) Quantification of cell cycle phase of tumor epithelial cells, grouped by patients. (E) UMAPs of tumor epithelial cells based on the top 4, 6, or 8 principal components, color-coded by patient, histological subtype, or the score of principal component 1; for patient color code see (G). (F) Heatmap of the top 30 genes positively or negatively correlated with principal component 1, respectively, shown for the top 500 cells with the highest or lowest PCA scores, respectively. (G+H) Distribution of (H) patients and (G) cell cycle phase of tumor epithelial cells sorted along principal component 1. (I) Mean module scores of the tumor cell signatures based on the top 30 genes positively or negatively correlated with principal component 1 (see (F)), respectively, in tumor epithelial cells sorted along principal component 1.

A

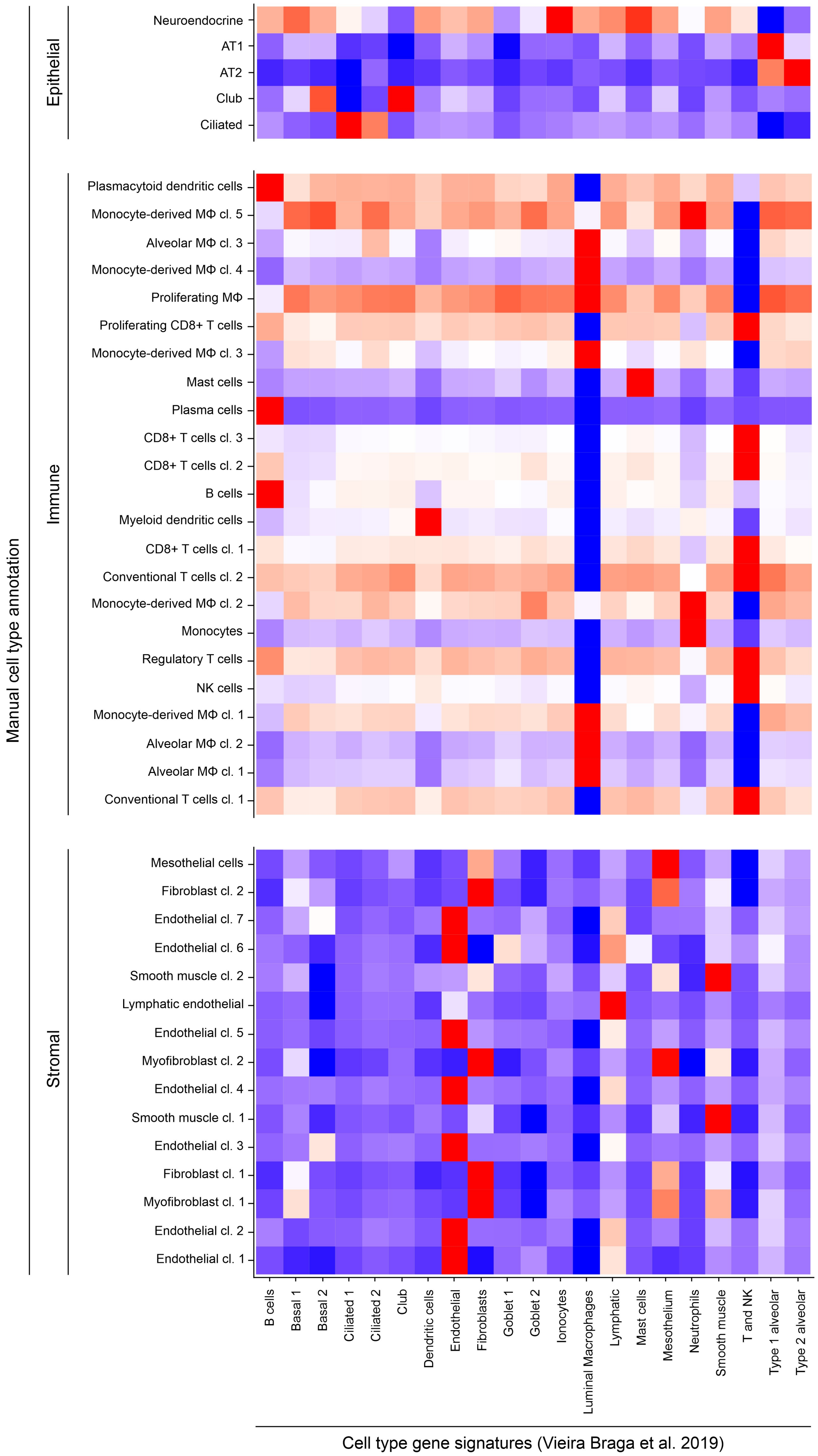

Manual cell type annotation
Epithelial
Neuroendocrine
AT1
AT2
Club
Ciliated
Immune
Plasmacytoid dendritic cells
Monocyte-derived MΦ cl. 5
Alveolar MΦ cl. 3
Monocyte-derived MΦ cl. 4
Proliferating MΦ
Proliferating CD8+ T cells
Monocyte-derived MΦ cl. 3
Mast cells
Plasma cells
CD8+ T cells cl. 3
CD8+ T cells cl. 2
B cells
Myeloid dendritic cells
CD8+ T cells cl. 1
Conventional T cells cl. 2
Monocyte-derived MΦ cl. 2
Monocytes
Regulatory T cells
NK cells
Monocyte-derived MΦ cl. 1
Alveolar MΦ cl. 2
Alveolar MΦ cl. 1
Conventional T cells cl. 1
Stromal
Mesothelial cells
Fibroblast cl. 2
Endothelial cl. 7
Endothelial cl. 6
Smooth muscle cl. 2
Lymphatic endothelial
Endothelial cl. 5
Myofibroblast cl. 2
Endothelial cl. 4
Smooth muscle cl. 1
Endothelial cl. 3
Fibroblast cl. 1
Myofibroblast cl. 1
Endothelial cl. 2
Endothelial cl. 1
B cells
Basal 1
Basal 2
Ciliated 1
Ciliated 2
Club
Dendritic cells
Endothelial
Fibroblasts
Goblet 1
Goblet 2
Ionocytes
Luminal Macrophages
Lymphatic
Mast cells
Mesothelium
Neutrophils
Smooth muscle
T and NK
Type 1 alveolar
Type 2 alveolar
Cell type gene signatures (Vieira Braga et al. 2019)

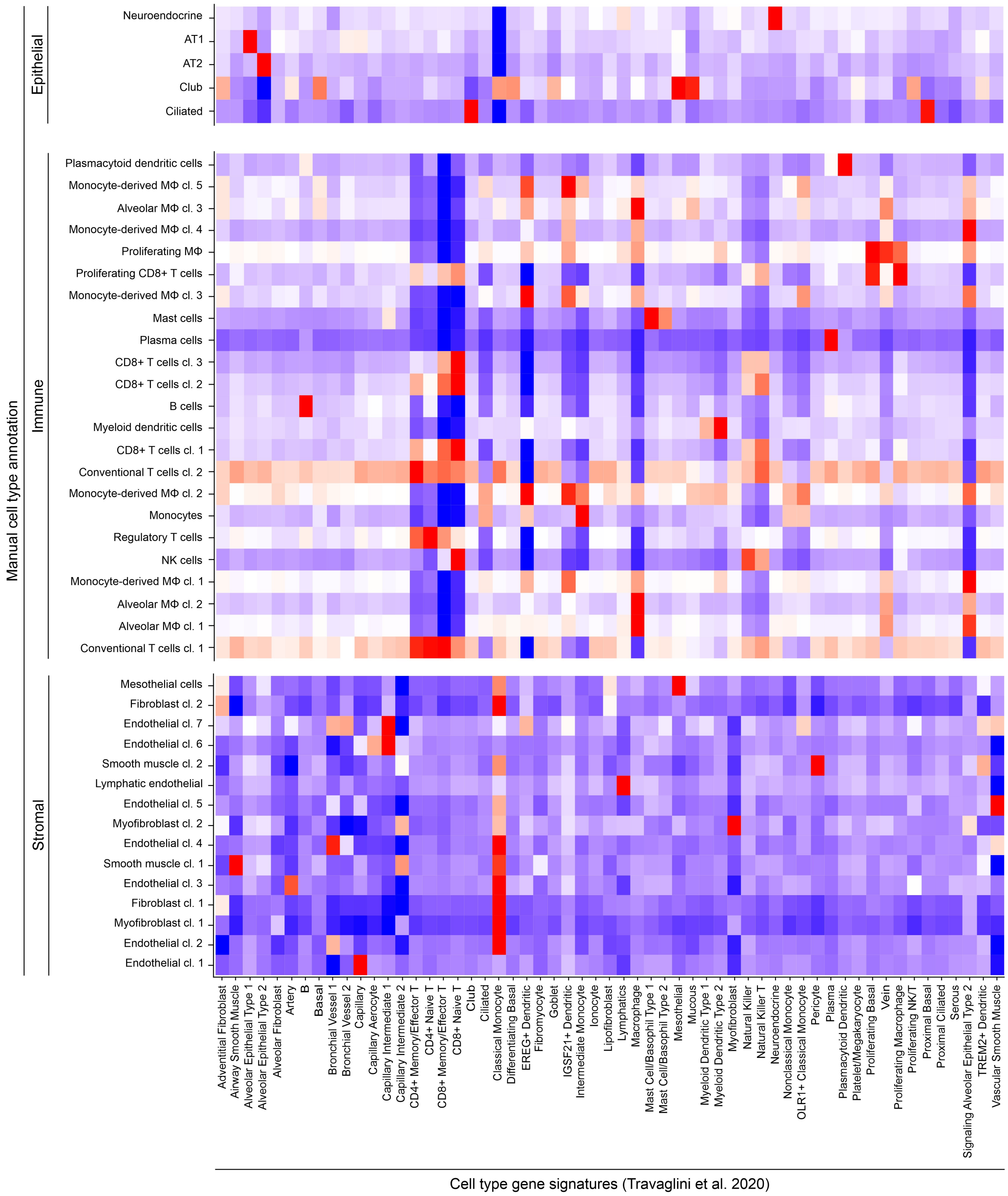


**Supplemental Figure 4**

(A+B) Mean module scores of cell type gene signatures according to (A) Vieira Braga et al. or (B) Travaglini et al. for manually annotated epithelial, immune and stromal cell clusters.

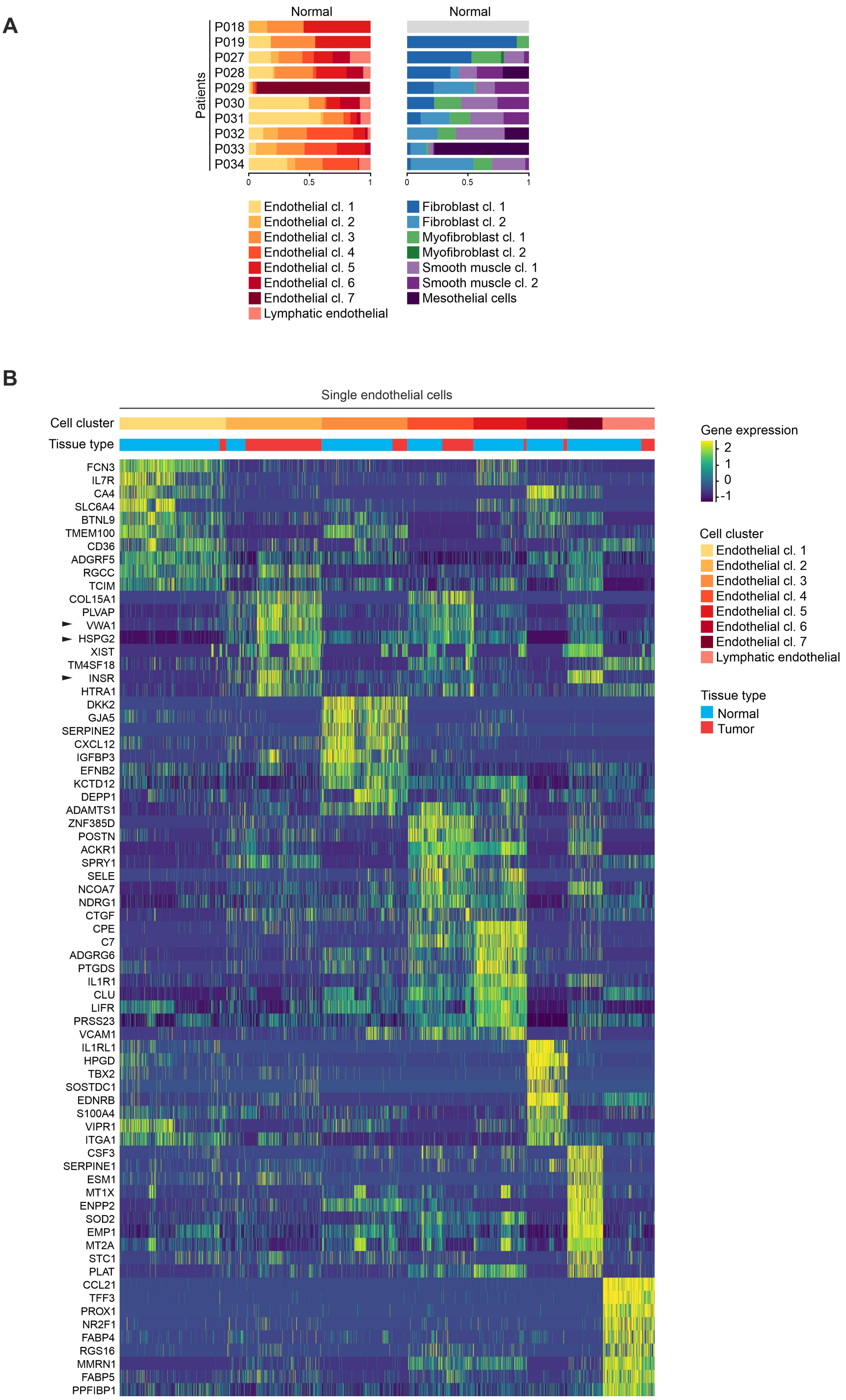


**Supplemental Figure 5**

(A) Quantification of endothelial and fibroblastic/muscle cell clusters per patient for normal tissue samples. (B) Differentially expressed genes of endothelial cell clusters, maximum top 10 genes showed per cluster, black arrowheads indicate genes mentioned in the main text.

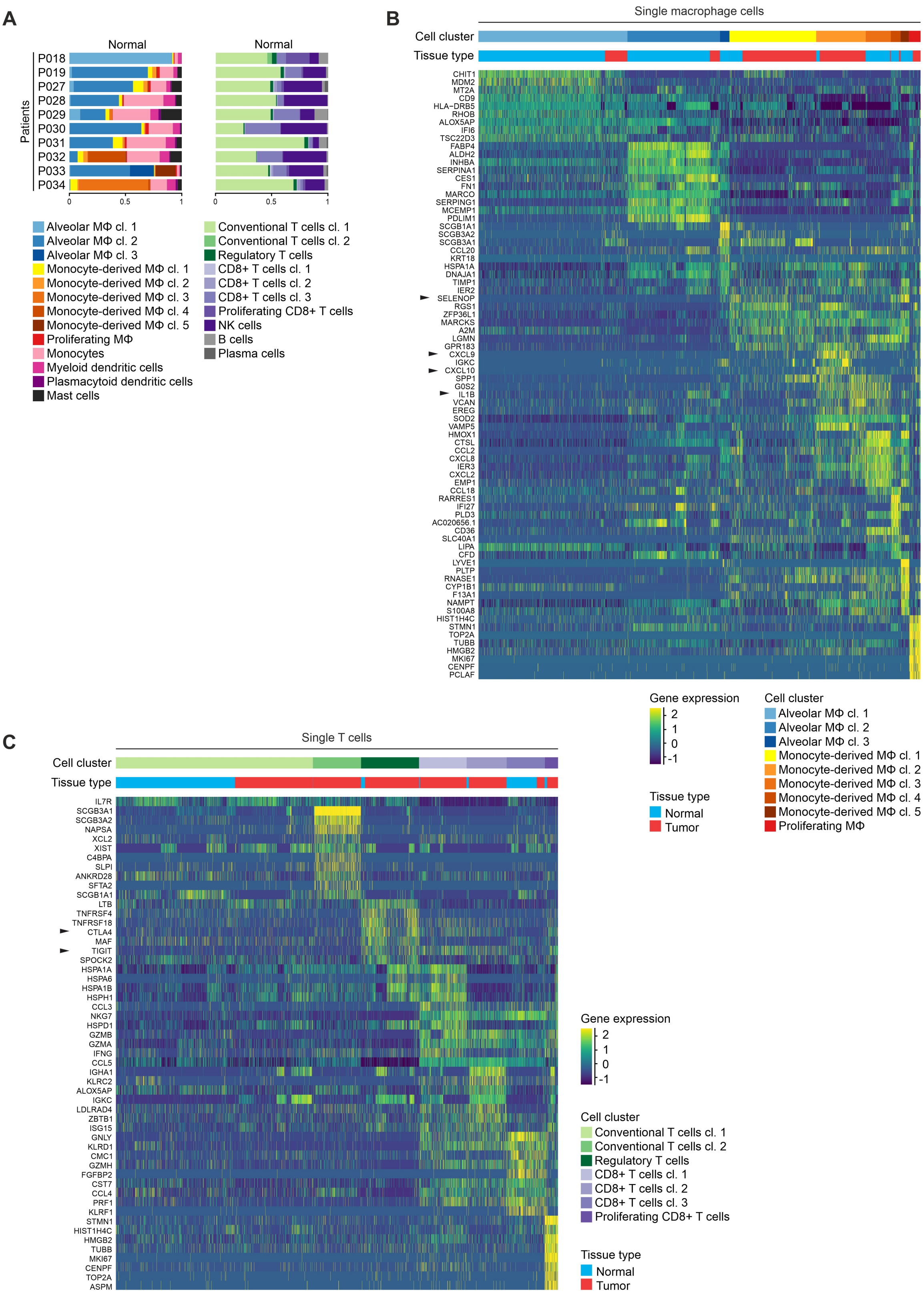


**Supplemental Figure 6**

(A) Quantification of myeloid and lymphoid cell clusters per patient for normal tissue samples. (B) Differentially expressed genes of macrophage cell clusters, maximum top 10 genes showed per cluster, black arrowheads indicate genes mentioned in the main text. (C) Differentially expressed genes of T cell clusters, maximum top 10 genes showed per cluster, black arrowheads indicate genes mentioned in the main text.

**A**

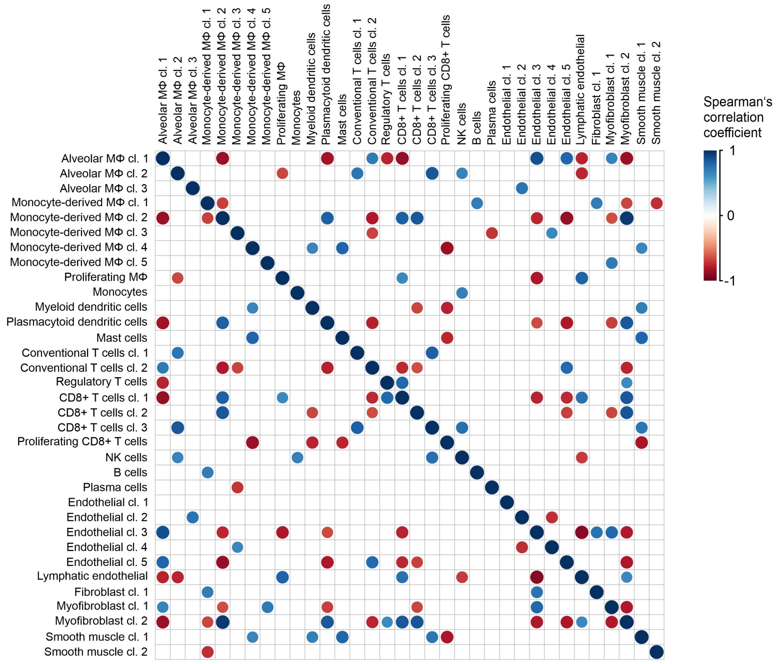


**B**

TCGA Lung adenocarcinoma

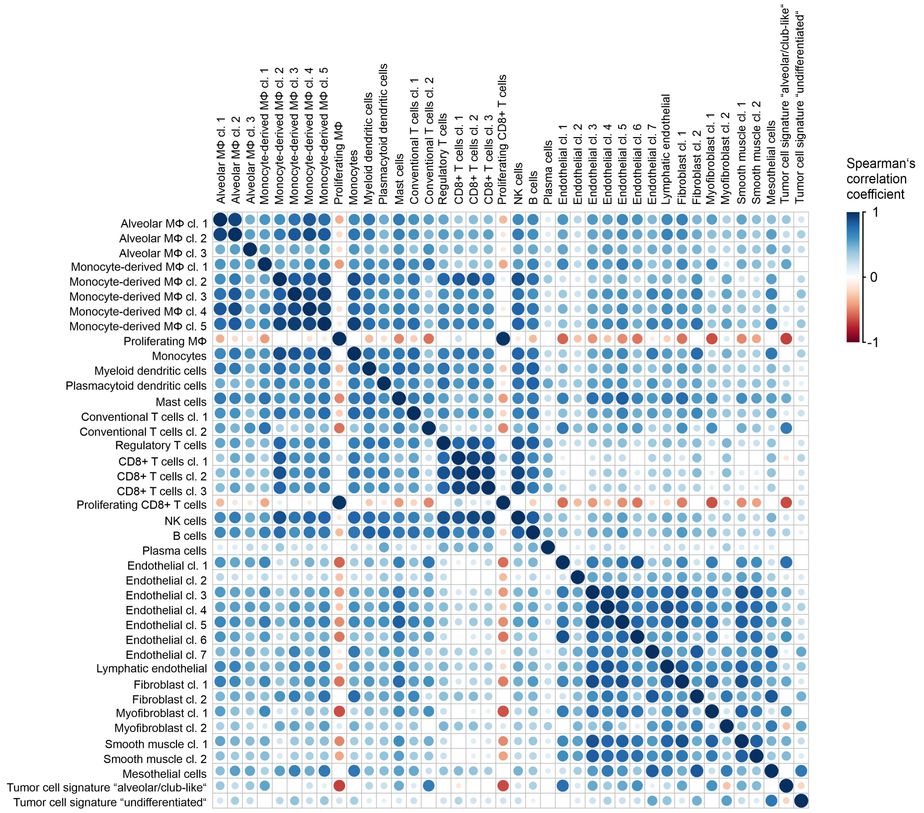


**Supplemental Figure 7**

(A) Correlation of the proportion of immune and stromal cell clusters in tumor tissue samples, cell clusters occurring in less than three tumor tissue samples were excluded from analysis; Spearman's correlation statistics, only correlations with $p < 0.05$ shown. (B) Correlation of ssGSEA enrichment scores of cell cluster marker gene signatures in the TCGA lung adenocarcinoma cohort; Spearman's correlation statistics, only correlations with $p < 0.05$ shown.
